# Pollutome complexity determines the removal of recalcitrant pharmaceuticals

**DOI:** 10.1101/2023.11.30.568980

**Authors:** Marcel Suleiman, Natalie Le Lay, Francesca Demaria, Boris A Kolvenbach, Mariana S Cretoiu, Owen L Petchey, Alexandre Jousset, Philippe F-X Corvini

## Abstract

Organic pollutants are an increasing threat for wildlife and humans. Managing their removal is however complicated by the difficulties in predicting degradation rates. In this work we demonstrate that the complexity of the pollutome, the set of co-existing contaminants, is a major driver of biodegradation. We built representative assemblages out of one to five common pharmaceuticals (caffeine, atenolol, paracetamol, ibuprofen, and enalapril) selected along a gradient of biodegradability. We followed their individual removal by wastewater microbial communities. The presence of multichemical background pollution was essential for the removal of recalcitrant molecules such as ibuprofen. Crucially, high order interactions between pollutants were a determinant, with the addition of new molecules particularly impacting assemblages of multiple compounds. We explain these interactions by shifts in the microbiome, with degradable molecules such as paracetamol enriching species and pathways involved in the removal of several organic molecules. We conclude that pollutants should be treated as part of a complex system, with emerging pollutants potentially showing cascading effects and offering leverage to promote bioremediation.

## Introduction

The widespread use of pharmaceuticals in society and agriculture as well as their unintentional release from production sites has led to an alarming increase in their presence and accumulation in wastewater treatment plants^1–3^. Organic pollutants pose a significant environmental concern due to their multiple and still poorly understood impacts on ecosystems and human health ^4,5^. Consequently, there is a growing need to understand and mitigate the impact of these diverse pollutants on biological processes in wastewater. Traditionally, research efforts have focused on studying the removal efficiency of individual/single pharmaceutical pollutants by microbial communities, which led to a classification of easily biodegradable, such as paracetamol ^6–8^, and recalcitrant micropollutants, such as ibuprofen and diclofenac ^8–12^. While these studies provided valuable insights into the degradation potential of specific compounds, they overlook the complexities of real-world scenarios, where several micropollutants co-occur ^13,14^. Here we assess whether the complexity of the pollutome, the set of nefarious molecules present in a given environment, can determine the removal of the pollutants present.

A growing number of studies have highlighted the unpredictable effects of multiple environmental pressures on ecosystems’ functioning, including microbes ^15,16^. Such unpredictable effects are also likely to happen within a pollutome: When microbial catabolic pathways overlap for specific pollutants, the enrichment of organisms degrading a compound may also promote the degradation of other harboring similar chemical patterns ^17–19^. In addition, induction of “promiscuous” enzymes with a large substrate spectrum may lead to broader pollutant removal ^20,21^. Further important mechanisms within a pollutome can probably be cross-feedings^22,23^, in which certain microorganisms form degradation metabolites to sustain other microorganisms’ growth, and co-metabolism ^24,25^, the transformation of a non-growth substrate in the obligate presence of a growth compound ^26^. To gain first insights into the impact of the presence of several pharmaceuticals on their removal in wastewater, we exposed wastewater samples to a combinatorial mixture of one to five commonly detected pharmaceuticals. We measured pollutant removal and bacterial community compositions. We hypothesize that due to pollutant-pollutant and pollutant-microbe interaction within the pollutome, recalcitrant pollutants can be more efficiently degraded.

## Methods

### Preparation of synthetic wastewater batch cultures

We set up a total of 96 batch cultures with a volume of 20 mL each (100 mL Erlenmeyer flasks). Each culture contained as a basis synthetic wastewater, following the OECD standard procedures (0.08 g/L peptone, 0.05 g/L meat extract, 15 mg/L urea, 3.5 mg/L NaCl, 2 mg/L CaCl2 x 2 H_2_O, 0.1 mg/L MgSO_4_ x 7 H_2_O and 1.4 mg/L K_2_HPO_4_, pH 7.5) (https://www.oecd.org/chemicalsafety/testing/43735667.pdf). Sterile filtered stock solutions of caffeine (C), atenolol (A), enalapril (E), paracetamol (P) and ibuprofen (I) were set up with a concentration of 10 g/L, and 200 µL were added to each batch culture, respectively, resulting in a final concentration of 100 mg/L of each pollutant. The five pollutants were added to the batch cultures in all possible combinations, leading to a total of 32 treatments (Table 1) (Supplementary Fig. 1 for chemical structures of the five pollutants). Each treatment was setup in triplicates.

**Table 1:**
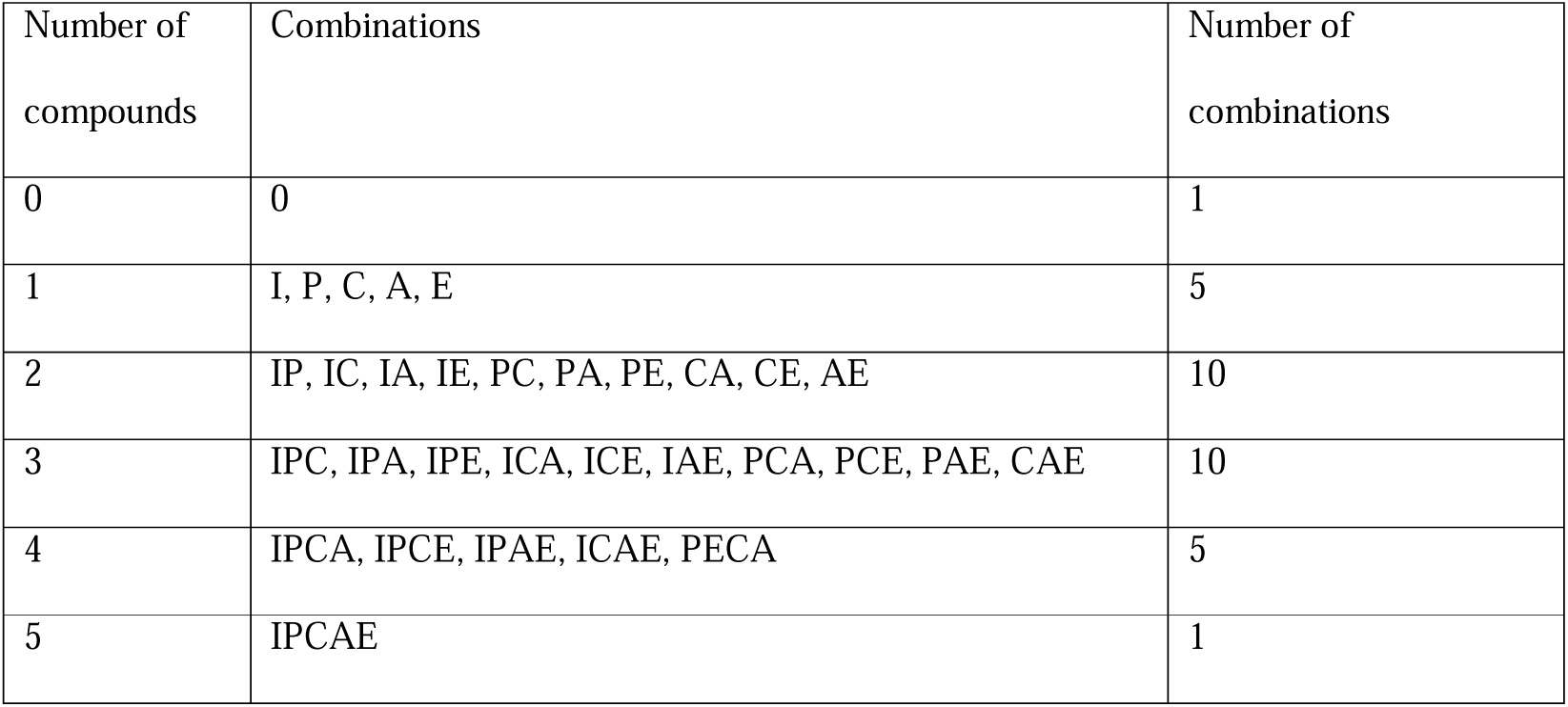
Summary of the combinations of pharmaceuticals in the batch cultures.

We used an additive design, meaning that each pollutant was added at a concentration of 100 mg/L. In addition, an abiotic control was set up in triplicate for each pollutant to ensure that no degradation occurred in absence of microorganisms. The batch cultures were inoculated with a sample from a wastewater treatment plant (Membrane bioreactor) with 1% (v/v). Incubation took place at 22°C for 11 days, under continuous shaking of the cultures (130 rpm).

### Sampling

On days 0, 3, 4, 7 and 11, one 500 µL sample was taken from each flask, centrifuged at 16,000 x g for 5 minutes and the filtered (0.45 µm pore filter) supernatant was used for HPLC analysis. In addition, on day 3 and day 11, 2 mL of culture was taken and centrifuged at 16,000 x g for 5 minutes, and the resulting pellet was taken for DNA extraction.

### HPLC analysis for measurements of pollutants

Pharmaceuticals were analyzed using high-performance liquid chromatography (HPLC) with a Hi-Plex Na column (Agilent Technologies). The separation was achieved by applying a flow rate of 0.7 mL/min, using a mobile phase consisting of a mixture of water and methanol. Detection of the pharmaceuticals was performed using a UV/VIS DAD detector. The initial mobile phase ratio was set at 80:20 VV, comprising 0.1% formic acid in Millipore water (A) and methanol (B). The B gradient was programmed to transition from 20% to 95% over a span of 15 minutes, enabling simultaneous analysis of all five micropollutants in a single run. The retention times for each pharmaceutical were as follows: ibuprofen eluted at 14.23 minutes, enalapril at 11.08 minutes, caffeine at 7.97 minutes, atenolol at 2.11 minutes, and paracetamol at 2.89 minutes. paracetamol, ibuprofen, atenolol and caffeine were detected at a wavelength of 230 nm, while enalapril was detected at 205 nm. Standard curves were generated for each pollutant, ranging from 1 mg/L to 500 mg/L (1 mg/L, 10 mg/L, 50 mg/L, 100 mg/L, 500 mg/L).

### DNA extraction and sequencing

DNA was isolated using the ZymoBIOMICS DNA Miniprep Kit (ZymoResearch, Irvine, USA) according to the manufacturer’s instructions. We sequenced the V4 region of the 16S rRNA gene (primer sequences 515f “GTGYCAGCMGCCGCGGTAA” and 806r “GGACTACNVGGGTWTCTAAT”) ^27^ using the Quick-16S™ Plus NGS Library Prep Kit (V4) (ZymoResearch) to create a DNA library. The library, containing 4 pM DNA (spiked with 25% PhiX), was sequenced in-house on an Illumina MiSeq platform, following the manufacturer’s instructions. Raw reads were processed using the R library *dada2* (Callahan et al., 2016). This involved quality control steps included analyzing primer sequences, assessing error rates (maxN=0, maxEE=c(2,2), truncQ=2), and identifying chimeras. The resulting sequence table (Min. number of reads = 187722, Max. number of reads = 2049193, Total number of reads = 160233034, Average number of reads = 88040) was aligned to the SILVA ribosomal RNA database (Quast et al., 2012) using version 138 (non-redundant dataset 99). A *phyloseq* object was then created using the *phyloseq* R library (McMurdie & Holmes, 2013).

This object consisted of an amplicon sequence variant (ASV) table, a taxonomy table, and sample data. For further analysis, the R libraries *phyloseq* (McMurdie & Holmes, 2013) and *vegan* ^28^ were employed. Functions of microbial communities were determined using PICRUSt2 ^29^. The *phyloseq* object, metadata, and detailed R code for analysis can be found on GitHub at https://github.com/Marcel2907. The raw sequencing data is available on the NCBI SRA (Sequence Read Archive) under the accession ID PRJNA1041291.

### Statistical analyses

A statistical model was used to analyze the degradation of each compound on each day. In each model the response variable was the percentage remaining of the focal compound on a specific day. In each model there were four binary explanatory variables, each of these coding the presence of the four non-focal compounds. All two-way and three-way interaction terms among explanatory variables were included, as well as the one four-way interaction. In all cases the model was a linear model with Gaussian errors (model diagnostics were acceptable). So, for example, with paracetamol as the focal compound, the model in *R* would be

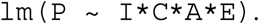

For the heatmaps in figure 1 and figure 3 (third column, respectively), the effect sizes for each day were calculated based on the summary of statistical analyses of pollutant interactions. Rows show the estimated coefficients of the single, one-way, two-way, three-way, and four-way interaction terms on pollutant concentration. White cells indicate a response variable and coefficient pairs for which the coefficients were not significantly different from zero (t-test p-value >.05), otherwise the diverging color palette illustrates the direction of the influence by the driver or interaction of drivers (based on the estimates of the *f*-test).

**Fig. 1.**
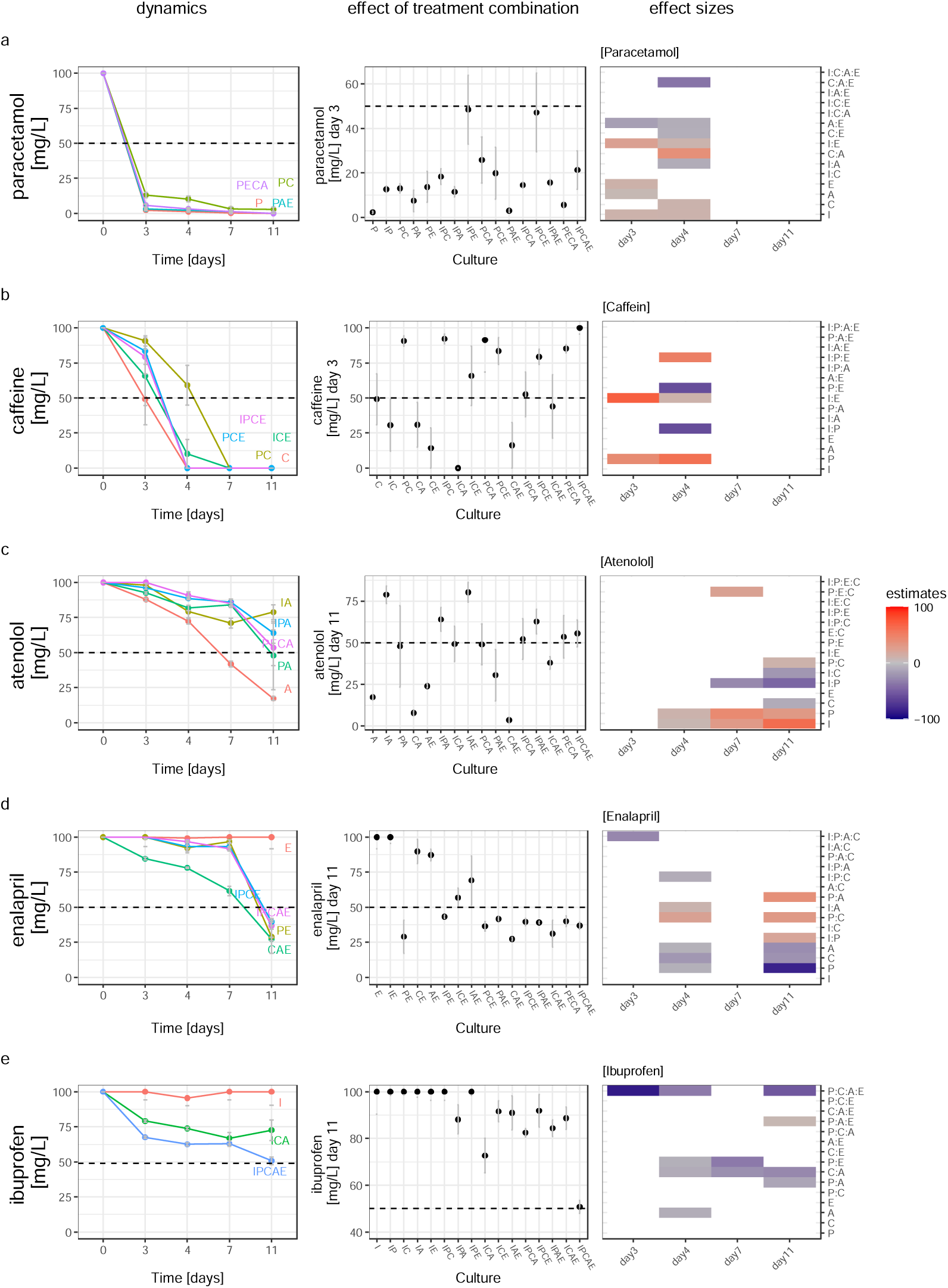
The presence of one or more pollutants increases the degradation of another pollutant. Each of the 5 pollutants appear in a row and the rows are sorted from least to most recalcitrant top to bottom. Dynamics and effect sizes of pharmaceuticals concentration within the batch cultures dependent on the presence of additional pharmaceuticals. Accentuation of the influence of pollutant combinations on the concentration of paracetamol (a), caffeine (b), atenolol (c), enalapril (d) and ibuprofen (e). Error bars show +/− SE (n=3). The effect sizes for each day are the summary of statistical analyses of pollutant concentrations (third column). Rows show the estimated coefficients of the single, one-way, two-way, three-way, and four-way interaction terms on pollutant concentration. White cells indicate a response variable and coefficient pairs for which the coefficients were not significantly different from zero (*t*-test *p*-value >.05), otherwise the diverging color palette illustrates the direction of the influence by the driver or interaction of drivers. Positive estimates are meaning higher concentrations of pollutants (i.e. lower degradation), while negative estimates are meaning lower concentrations of pollutants (i.e. higher degradation). A: Atenolol, C:Caffein, E: Enalapril, I: Ibuprofen, P: Paracetamol.

All statistical analyses can be found in detail in supplementary table 1 and 2.

## Results

### Removal kinetics of single and multiple pharmaceuticals in batch cultures

Degradation rates of single pharmaceutical substances in isolation clustered them in two groups: Category 1 (degradable) encompassed caffeine, paracetamol, and atenolol. Caffeine and paracetamol were completely removed within 3-4 days, while about 80% of atenolol was removed within 11 days (Fig. 1a,b,c; Supplementary Fig. 2). The second category (recalcitrant) contained enalapril and ibuprofen, which were not degraded when individually present (Fig 1d, e; Supplementary Fig. 2).

The degradation rates in mixed pharmaceuticals strongly departed from this baseline. The presence of degradable pharmaceuticals increased the degradation of ibuprofen and enalapril. For example, ibuprofen concentration was reduced to 50% of the original concentration when present alongside all four other pharmaceuticals, and to 70-90% of the initial concentration in some combinations of two or three other compounds (Fig. 1e). Enalapril was reduced to approximately 30% of its initial concentration when paracetamol was also present and or when paracetamol was absent but both atenolol and caffein (“CAE”) were present (Fig. 1d).

While pollutants of category 1 were to some extent degradable in all batch cultures, some inhibition effects were observed. For instance, atenolol degradation was hindered in the presence of ibuprofen or paracetamol, and by the presence of various other combinations of other compounds (Fig. 1c). In contrast, no striking additive negative effect was observed when atenolol was present alongside ibuprofen and paracetamol together (Fig. 1c “IPA”). Also, the removal of caffeine was slower in the presence of other pharmaceuticals, especially when paracetamol was present a long lag-phase occurred (Fig. 1b). In contrast, paracetamol degradation was only marginally inhibited by the presence of caffeine (Fig. 1a).

These effects of the presence of combinations of other pharmaceuticals on the degradation of a pharmaceutical were not only visually clear but were also clearly revealed by statistical analyses (Fig. 1 third column, Supplementary Table 1). These analyses revealed strong evidence of 2-way, 3-way, and 4-way interaction effects and dependencies among pollutants in the removal processes for all tested pharmaceuticals and were statistically confirmed.

The findings suggest that introducing one/more specific pollutant/s significantly affects the degradation of another compound, with the outcome being influenced by the presence of other pollutants. Particularly in the case of ibuprofen removal, identified as the most recalcitrant substrate in this study, notable variations were observed depending on the presence or absence of enalapril. This made up to 30% difference (Fig. 2).

**Fig. 2.**
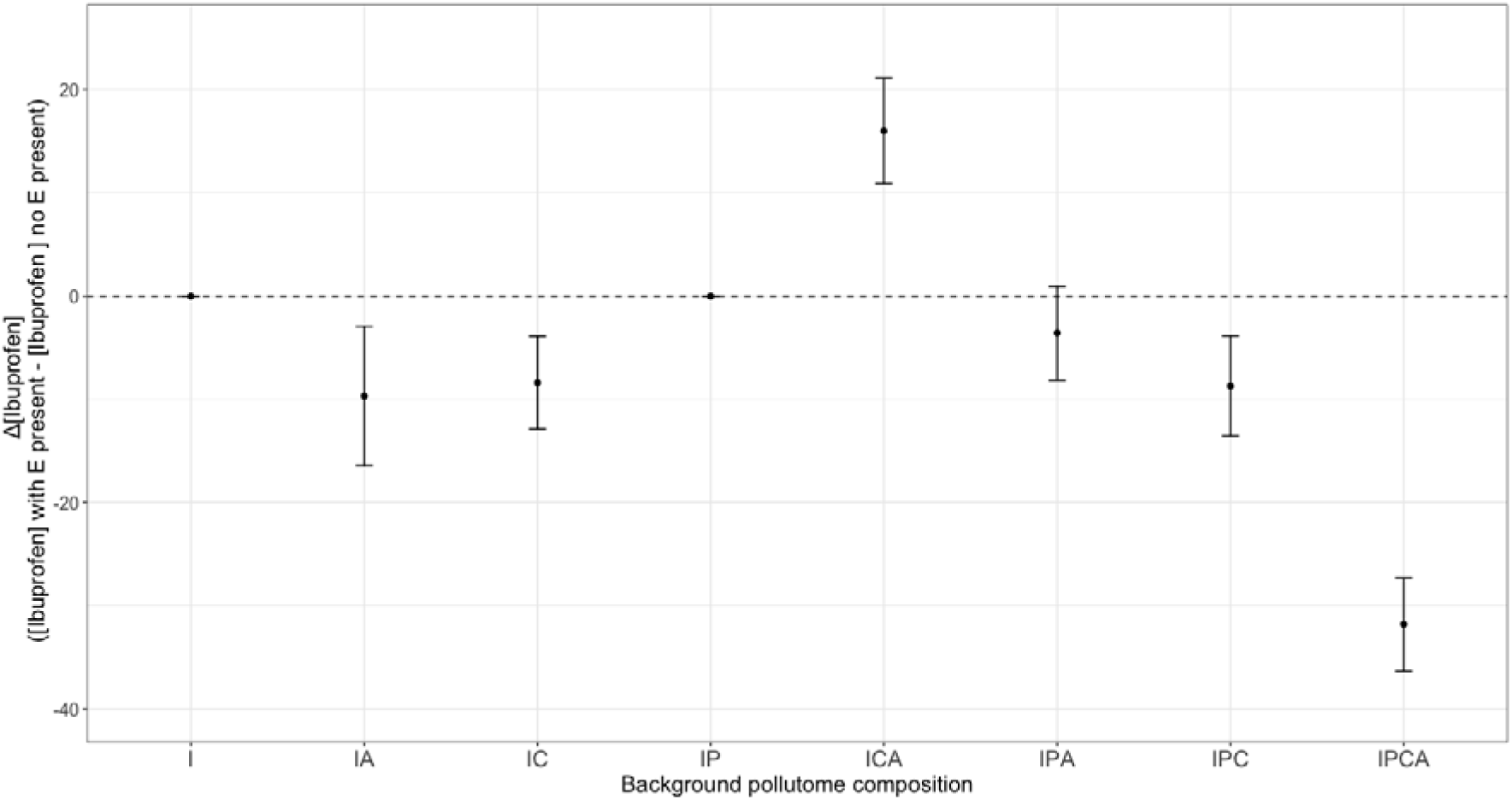
Pollutome complexity impacts the effect of additional pollutants on the biodegradation process. This figure shows the effect of enalapril addition on the biodegradation rate of the recalcitrant pollutant ibuprofen. Pollutome complexity is defined here as the number and identity of pharmaceuticals in the mixture. Impact of enalapril is defined as the average difference in ibuprofen concentration in a mix of pollutant with or without enalapril. Enalapril addition had little effect on the biodegradation of ibuprofen alone (I), but strongly increased its removal when all other pollutants were present (IPCA). A: Atenolol, C:Caffein, E: Enalapril, I: Ibuprofen, P: Paracetamol. Error bars show +/− SE (n=3).

### Microbial community composition dependencies on pharmaceutical combinations

Biomass increased in all batch cultivations, indicating an active microbial community. On day 3, biomass was significantly higher in the presence of single degradable pollutants (atenolol, caffeine, paracetamol) compared to pharmaceutical-free controls or single recalcitrant pollutants (ibuprofen, enalapril) (Figure 3a, main effects in Supplementary Table 2). There was no clear relationship between the number of pollutants and microbial biomass (*f*-test *p*-value 0.57 (number of stressors as a factor)). There is perhaps a pattern of higher biomass when certain combinations of pollutants are present (i.e., PA, PCA, IPCA, ICAE, PECA, and IPCAE). Furthermore, while ibuprofen had uniquely high degradation in the IPCAE treatment, that treatment did not have uniquely high microbial biomass (e.g., the background community ICAE without or with P had similar microbial biomass, statistics in Supplementary Table 2), suggesting at least for ibuprofen that differences in microbial biomass did not drive the observed difference in degradation. Depending on the combination of pharmaceuticals, the biomass mostly decreased from day 3 to day 11, with some exceptions (Fig. 3a, “C”, “ICA”).

**Fig. 3.**
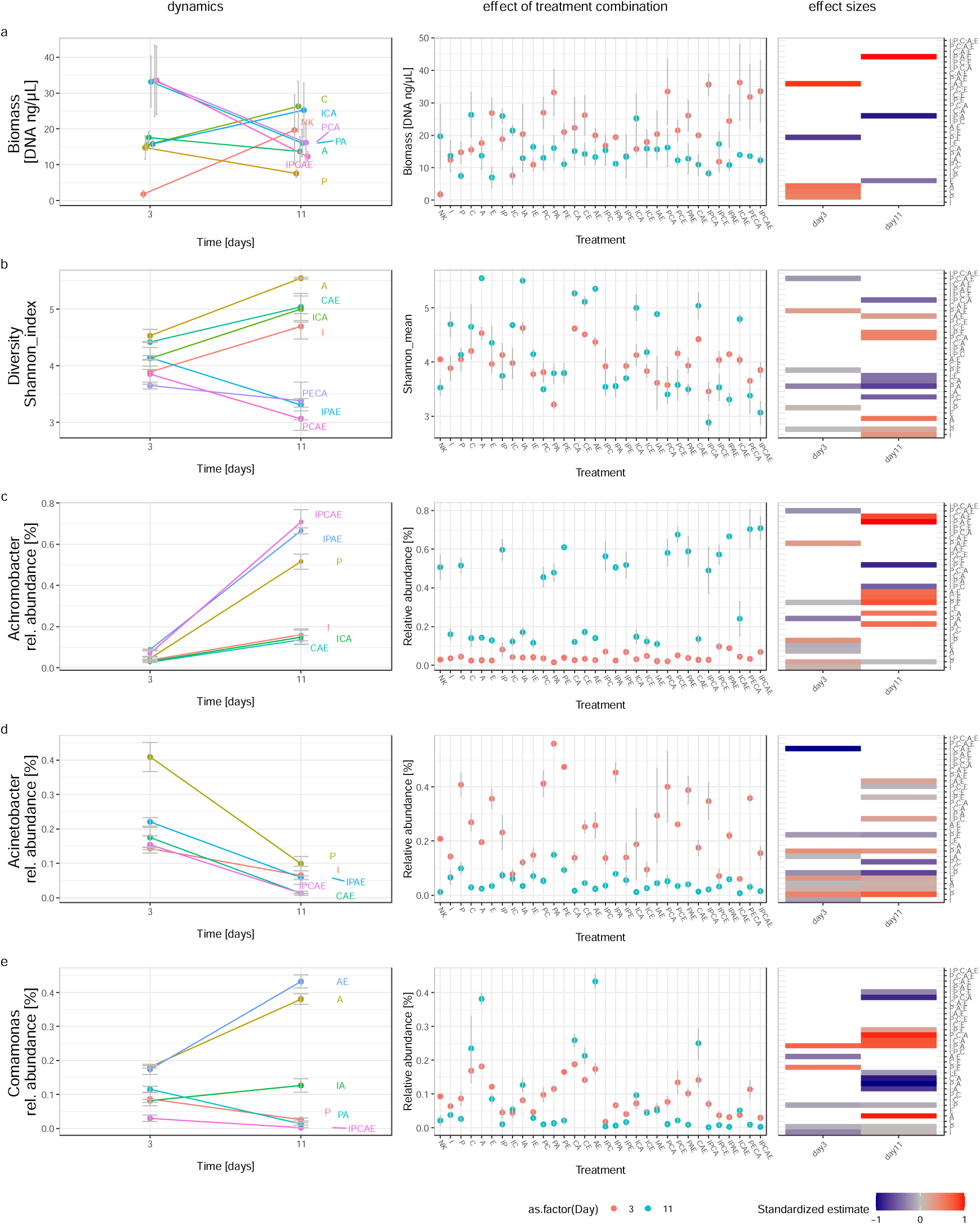
Dynamics and effect sizes of microbial community variables dependent on pharmaceutical combinations. Dynamics and effect sizes are showing the influence of pollutant combination on biomass (a), diversity (b), relative abundance of *Achromobacter* (c), *Acinetobacter* (d) and *Comamonas* (e). The effect sizes for each day are the summary of statistical analyses of pollutant concentrations (third column). Rows show the estimated coefficients of the single, one-way, two-way, three-way, four-way and five-way interaction terms on pollutant concentration. White cells indicate a response variable and coefficient pairs for which the coefficients were not significantly different from zero (*t*-test *p*-value >.05), otherwise the diverging color palette illustrates the direction of the influence by the driver or interaction of drivers (estimates of each variable were standardized by dividing by the largest absolute value of the estimates in each variable). NK = control (synthetic wastewater without addition of pharmaceuticals). A: Atenolol, C:Caffein, E: Enalapril, I: Ibuprofen, P: Paracetamol.

Shannon index, the reference index used to depict biodiversity, was strongly impacted by pharmaceutical treatment combinations (Fig. 3b). Further, the presence of atenolol increased Shannon index in all cultures, regardless of the number of pharmaceuticals present (main effect of atenolol Supplementary Table 2).

A linear model was also used to analyze how microbial biomass, microbial diversity, and the relative abundance of three most abundant taxa depended on the five compounds and their interactions.

The microbial communities showed strong differences depending on the pharmaceutical treatment on day 3 and day 11. On day 3, genera with high relative abundance within the microbial communities were identified as members of the genera *Achromobacter*, *Trichococcus*, *Acinetobacter*, *Pseudomonas, Comamonas,* whose abundance strongly varied in different treatments (Fig 3c,d,e, Supplementary Fig. 4). For example, *Trichococcus* showed a relative abundance of 17-27% in triplicates incubated with PECA but was below detection when cultivated with enalapril (E) only. Interestingly, comparable treatment effects were observed for these genera on day 11, but in different strengths (Fig. 3c,d,e, second column). Additionally on day 11, the proportion of members of the genus *Achromobacter* strongly increased. This was also highly influenced in their magnitude dependent on the treatment combination (Fig. 3c, Supplementary Fig. 3). *Achromobacter* abundance was consistently increased by the presence of paracetamol, an effect reflected by non-metric multidimensional scaling analysis (NMDS), which showed a strong clustering of paracetamol cultures on day 11 (Fig. 4a). Interestingly, on day 11, paracetamol was already degraded (within 3 days). Therefore, *Achromobacter* may possibly be associated with the consumption of degradation products of paracetamol like aminophenol which occurrence was verified by HPLC. Statistical analysis confirmed significant changes of relative species abundance caused by different combinations of pharmaceuticals (Fig. 3c,d,e, third column). *Achromobacter* abundance also increased in the negative control (no pollutant added), but NMDS indicates a different community composition in these controls compared to paracetamol-containing cultures (Fig. 4a,b, Supplementary Fig. 3, Supplementary Table 2).

**Fig. 4.**
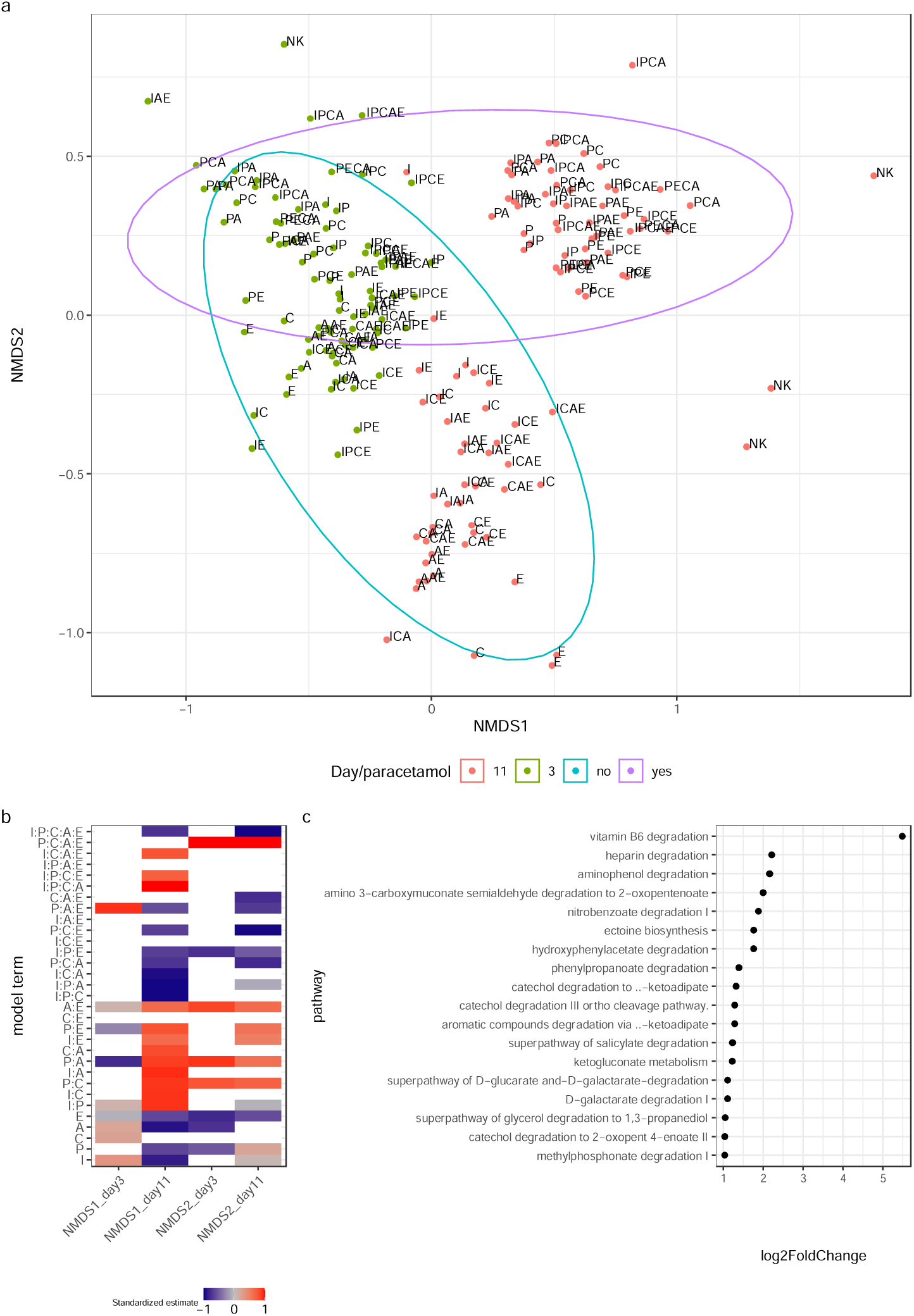
Effect of pharmaceutical combinations on microbial community structure and function. (a) Beta-diversity based on NMDS analysis (Bray-Curtis) in relation to the number of micropollutants present and day. All samples containing paracetamol on day 11 are clustering on the top right. Stress is 0.17. (b) The effect sizes for each day are the summary of statistical analyses of pollutant concentrations for NMDS1 and NMDS2. Rows show the estimated coefficients of the single, one-way, two-way, three-way, and four-way interaction terms on pollutant concentration. White cells indicate a response variable and coefficient pairs for which the coefficients were not significantly different from zero (*t*-test *p*-value >.05), otherwise the diverging color palette illustrates the direction of the influence by the driver or interaction of drivers (estimates of each variable were standardized by dividing by the largest absolute value of the estimates in each variable). NK = control, no pollutant added (just synthetic wastewater). (c) Significant (p > 0.01) over-represented pathways of microbial communities grown on paracetamol (also in all combinations with other pollutants) versus non-paracetamol containing batch cultures. Analysis was performed using picrust2^29^ and deseq2 ^30^. Pathways were taken from Metacyc^31^. The most significant affected pathways are shown. A: Atenolol, C:Caffein, E: Enalapril, I: Ibuprofen, P: Paracetamol.

Since microbial communities incubated with paracetamol exhibited a different community than all other treatments (day 11; Fig. 4a), we compared their inferred functional gene profiles (Fig. 4c) using PICRUSt2 ^29^. Paracetamol significantly enriched pathways for aminophenol degradation, catechol degradation, as well as several aromatic degradation pathways (Fig. 4). As these pathways are likely involved in the degradation of a broad range of organic molecules, their increase may explain the positive effect of paracetamol on other more recalcitrant pharmaceuticals.

## Discussion

Efficient pollutant removal from wastewater is essential for environmental safety, yet current water treatment facilities fail to remove organic pollutants such as pharmaceuticals ^32^. Steering microbial communities within these unique ecosystems may be key to designing better removal strategies. Microbial community dynamics can rapidly change in composition and function depending on the incoming water composition^33^. Therefore, these wastewater treatment plants can be seen as a model system for studying multiple drivers on microbial communities and their degradation capacity of pollutants. Pollutant removal has been extensively studied in isolation, providing detailed insights into the molecular mechanisms underlying biodegradation. However, these findings only marginally translate to real-world scenario, where multiple drivers co-occur ^34,35^.

In this work we shed light on the interactive effects of pollutants within the pollutome, using mixtures of pharmaceutical varying in biodegradability as a model. We demonstrate that the complexity of the pollutome is a major driver of biodegradation and that the presence of multichemical background pollution was essential for the removal of recalcitrant molecules. We found in particular the degradation of recalcitrant pollutants to be strongly modulated by the presence of other pollutants. Easily degradable pharmaceuticals such as paracetamol, atenolol, and caffeine enable the degradation of the more recalcitrant ibuprofen and enalapril. In addition, some pollutants may hinder the biodegradation of other. This was particularly striking for atenolol, which degradation was notably inhibited in the presence of ibuprofen. These two chemicals show strong structural similarities (*e.g.* benzene ring and alkyl chain), which might inhibit enzymatic activity. Ibuprofen deserves special attention, as it proved to be only degradable when incubated alongside other pharmaceutical compounds. This observation underscores the significance of studying the environmental fate of pharmaceuticals as a collective group rather than at single compound level.

Interactions between pollutants are likely due to shifts in microbial community composition and function. We identified a range of potential key players (based on high relative abundances) for pharmaceuticals, namely *Achromobacter*, *Pseudomonas*, *Acinetobacter*, *Comamonas* and *Trichococcus*. All of them have already been shown to be associated with pollutant degradation ^6,7,36–39^. These genera strongly respond to the composition of the pollutant mix, potentially explaining previous observations of their fluctuations in wastewater treatment systems ^40,41^. In particular, paracetamol had a strong effect on the microbial community composition. When degraded, paracetamol is broken down into aminophenol, which is toxic for many microorganisms ^42^. Paracetamol-treated communities indeed showed an increased abundance of the aminophenol pathway. This pathway may have served as detoxification mechanism ^7^ but may also be involved in the degradation of other recalcitrant pharmaceuticals. It’s also possible that the pathways involved in paracetamol degradation could lead to the breakdown of another substance, utilizing enzymes such as monooxygenase and dioxygenase, for instance. One caveat to mention here is the fact that due to the comparatively high concentration used in this study, it is possible that the microbial community was not able to degrade fast enough the formed aminophenol, which might have accumulated transiently.

The major question which needs to be tackled in future studies is the reason for this observation. In general, underlying reason could be rooted in mechanisms like cross-feedings, elevated enzyme activity, increased energy levels, and the induced expression of genes encoding promiscuous enzymes^20–23,26^. However, our study can probably rule out increased biomass as a contributing factor since the ibuprofen-degrading cultures did not yield more biomass than other treatments. Co-metabolic effect related to enzyme that fortuitously accept various chemically related substrate could play a role, since the chemical structures of the used pharmaceuticals show some chemical similarities, such as aromatic rings (Supplementary Figure 1). Therefore, it could be possible that e.g. specific dioxygenases that play a role in paracetamol degradation ^7,43^ could potentially also show (low) activity against ibuprofen and enalapril.

All tested pharmaceuticals are globally detectable in wastewater influents and effluents and occur worldwide in ng/L -µg/L scale ^6,44–47^, which is significantly lower compared to the high concentration we used in this study. However, the CODs used in this study can occur in wastewater of industrial production sites of pharmaceuticals ^48^.

This study indicates that the presence of easily degradable micropollutants, such as caffeine, atenolol, and paracetamol, promoted the degradation of recalcitrant substrates like ibuprofen and enalapril. In contrast, these latter compounds were not degraded when present as the sole pollutant. The significance of these discoveries is noteworthy, as they can serve as potential starting points for the development of future applications aimed at the effective removal of pharmaceuticals: The study demonstrated that the addition of specific compounds at specific time points can enhance degradation of a target pollutant. Addition of non-toxic functional mimics of existing pollutants may thus improve the microbial removal of persistent pollutants, contributing to safe water, ecosystems, and food supply. We conclude that pollutants should be treated as part of a complex system, with emerging pollutants potentially showing cascading effects and offering leverage to promote bioremediation.

## Declarations

## Data availability

All datasets and metadata are available on the GitHub repository Marcel29071989 (https://github.com/Marcel29071989/), and the raw sequencing data can be found on NCBI SRA archive under ID PRJNA1041291.

## Code availability

R Studio code for the analysis and plotting of figures for the manuscript and supplementary information is available at https://github.com/Marcel29071989

## Competing interests

The authors declare that they have no competing interests.

## Funding

This work was funded through the European Union’s Horizon 2020 project NMYPHE under grant ID 10106625.

## Authors’ contributions

Marcel Suleiman and Philippe Corvini planned the experimental setup. Natalie La Ley performed all experiments in the lab. Francesca Demaria and Boris Kolvenbach supported analytical lab work and Boris Kolvenbach supported the analysis of metabolic pathways. Marcel Suleiman performed the up- and downstream bioinformatics. Marcel Suleiman, Owen L. Petchey and Alexandre Jousset performed the data analysis. Owen Petchey and Alexandre Jousset supported the use of statistical models. Silvia Cretoiu performed the Picrust2 analysis and gave bioinformatic support. Owen L. Petchey, Marcel Suleiman, Philippe Corvini and Alexandre Jousset drafted the manuscript. All authors confirmed the final version of the manuscript.

## Acknowledgements

Not applicable

## Ethics approval and consent to participate

Not applicable

## Consent for publication

Not applicable

## Notes

### Competing Interest Statement

The authors have declared no competing interest.

