## Supplementary Information for "Pollutome complexity determines the removal of recalcitrant pharmaceuticals"

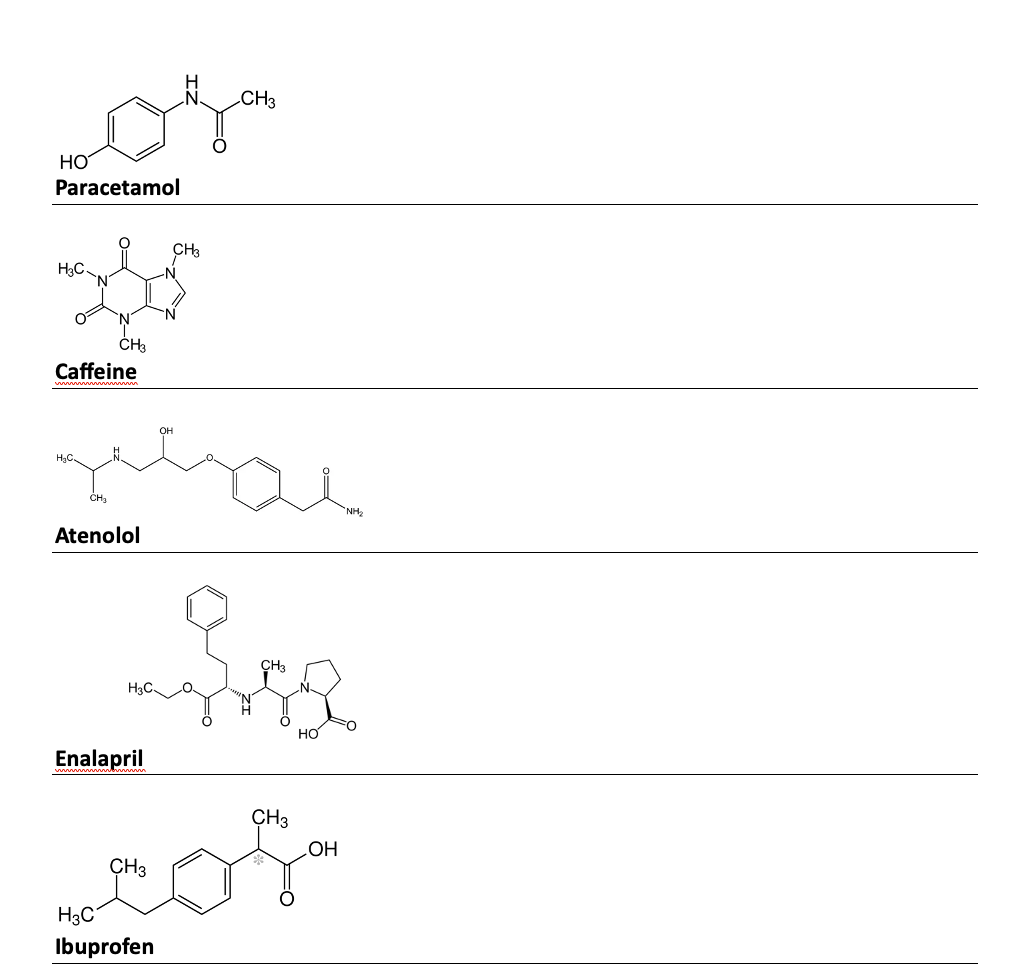

Supplementary Fig. 1 Chemical structure of the five pollutants of this study.

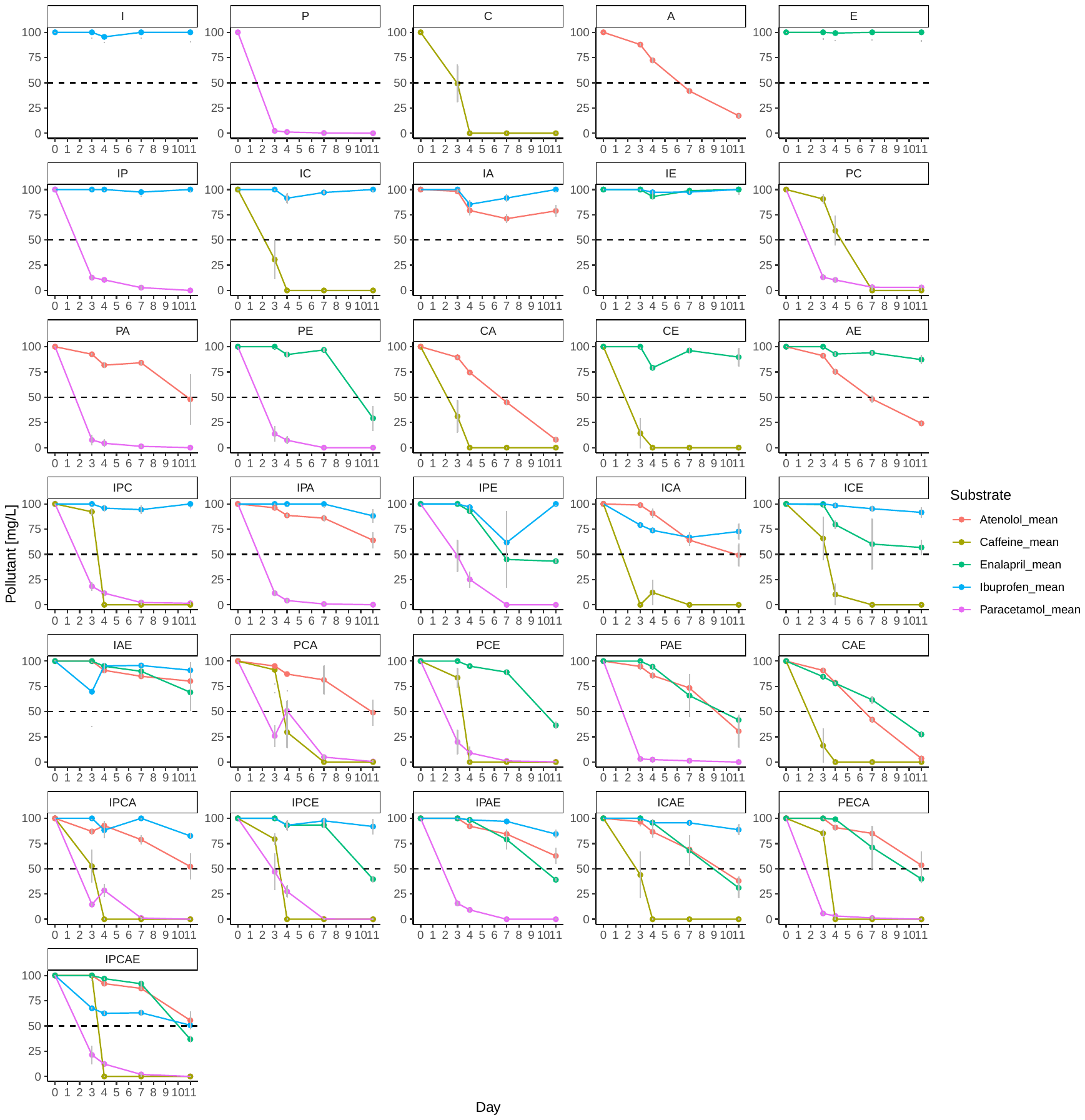

Supplementary Fig.2 Concentration (in mg/L) of five pharmaceuticals in batch cultures. Time series of the concentration (in mg/L) of atenolol, caffeine, paracetamol, enalapril and ibuprofen in all batch cultures incubated for 11 days in all possible combinations. Error bars are shown (n=3). I: Ibuprofen; P: Paracetamol; C: Caffeine; A: Atenolol; E: Enalapril

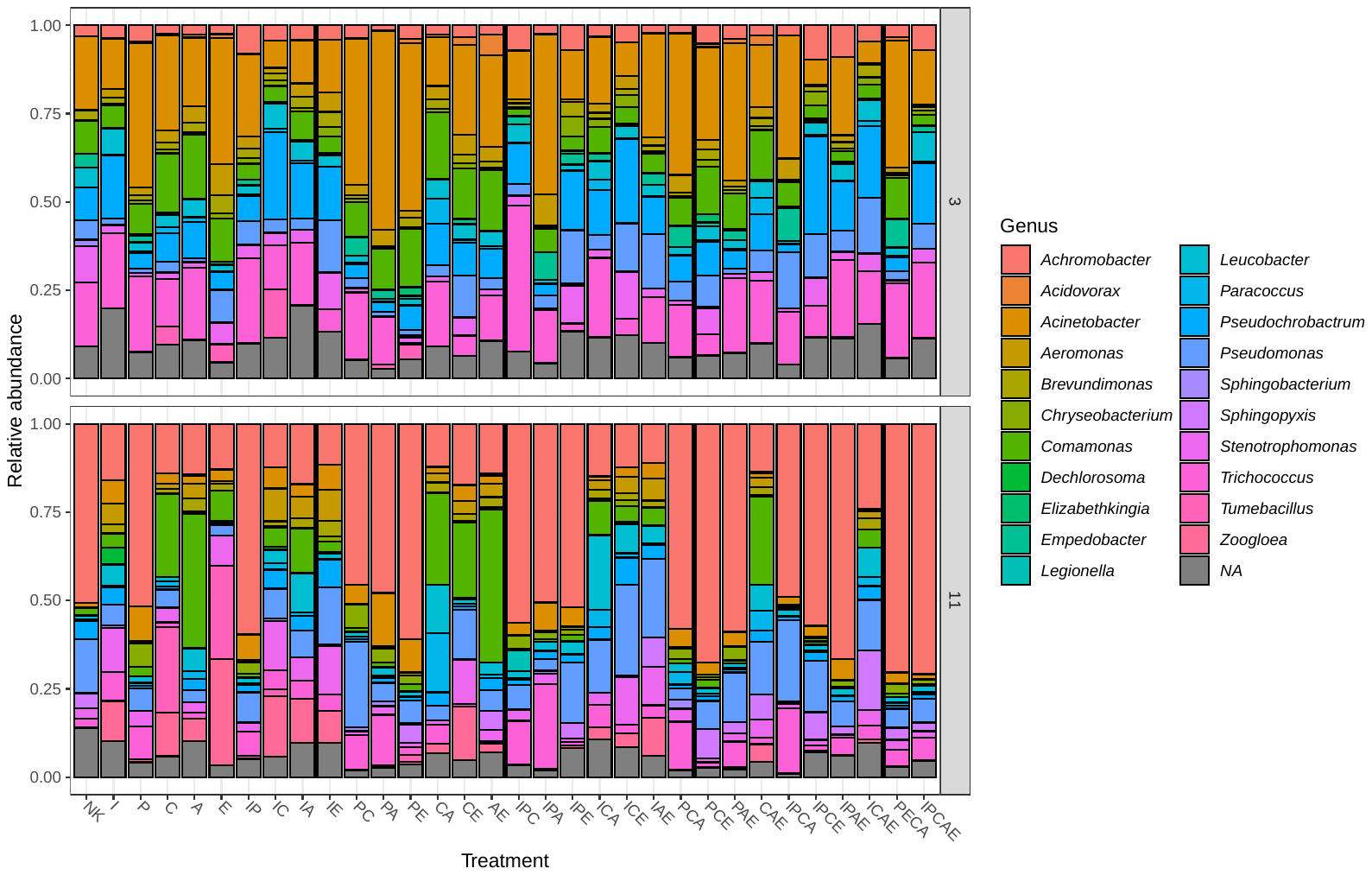

Supplementary Fig.3 Relative community composition (in %) of genera in dependence of the micropollutant present on day 3 and day 11. Just genera with a relative abundance of < 4 % in at least one sample are shown. Each treatment is represented by the mean of three replicates. A: Atenolol, C:Caffein, E: Enalapril, I: Ibuprofen, P: Paracetamol.

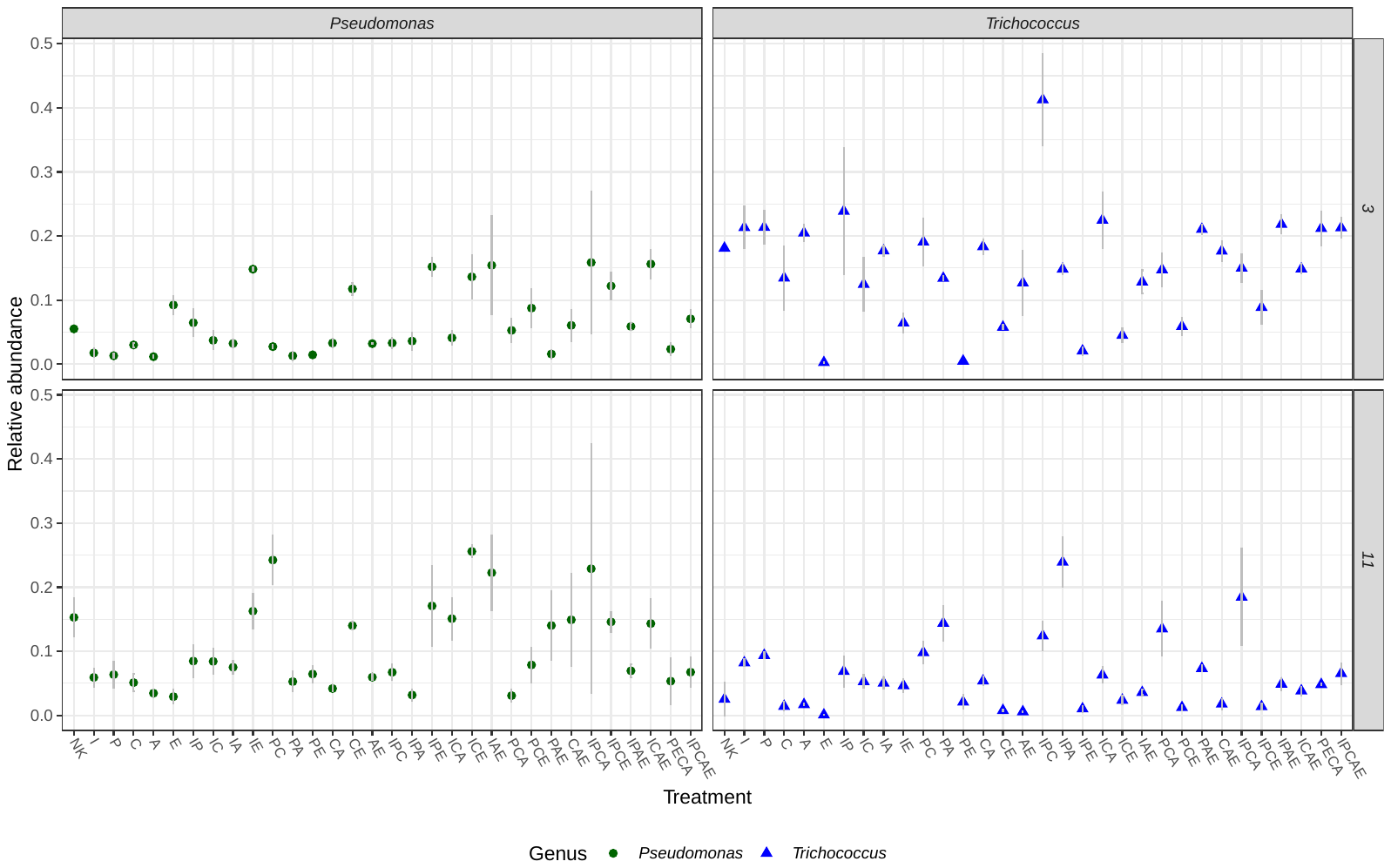

Supplementary Fig. 4 Influence of pharmaceuticals combination on the relative abundance of *Pseudomonas* and *Trichococcus*. Relative abundance (in %) of five most abundant microbial genera on day 3 and day 11 in dependence on the micropollutant combination. Symbols are the mean of the three replicates, and errors bars show the standard error of the mean (n=3). A: Atenolol, C:Caffein, E: Enalapril, I: Ibuprofen, P: Paracetamol.

Supplementary Table 1: Statistic analysis of the pollutant concentration based on pharmaceutical combination. A separate statistical model was used to analyze the degradation of each compound on each day. In each model the response variable was the percentage remaining of the focal compound on a specific day. In each model there were four binary explanatory variables, each of these coding the presence of the four non-focal compounds. All two-way and three-way interaction terms among explanatory variables were included, as well as the one four-way interaction. In all cases the model was a linear model with Gaussian errors (model diagnostics were acceptable). So, for example, with paracetamol as the focal compound, the model in *R* would be lm(P ~ I*C*A*E).

| **treatment_combination** | **estimate** | **std.error** | **t_statistic** | **t_p.value** | **df** | **sumsq** | **meansq** | **f_statistic** | **f_p.value** | **variable** | **compound** |
| --- | --- | --- | --- | --- | --- | --- | --- | --- | --- | --- | --- |
| P | 2.053038 | 11.66958 | 0.175931 | 0.861457 | 1 | 107.8474 | 107.8474 | 0.527968 | 0.472744 | day3 | [Ibuprofen] |
| C | 2.053038 | 11.66958 | 0.175931 | 0.861457 | 1 | 68.21605 | 68.21605 | 0.333952 | 0.567386 | day3 | [Ibuprofen] |
| A | 1.93718 | 11.66958 | 0.166003 | 0.869199 | 1 | 1362.177 | 1362.177 | 6.668549 | 0.014593 | day3 | [Ibuprofen] |
| E | 2.053038 | 11.66958 | 0.175931 | 0.861457 | 1 | 341.6687 | 341.6687 | 1.672642 | 0.205164 | day3 | [Ibuprofen] |
| P:C | -2.05304 | 16.50328 | -0.1244 | 0.901775 | 1 | 401.6576 | 401.6576 | 1.966319 | 0.170466 | day3 | [Ibuprofen] |
| P:A | -1.93718 | 16.50328 | -0.11738 | 0.907291 | 1 | 74.08008 | 74.08008 | 0.36266 | 0.551276 | day3 | [Ibuprofen] |
| C:A | -22.8876 | 16.50328 | -1.38685 | 0.175073 | 1 | 100.7471 | 100.7471 | 0.493208 | 0.487577 | day3 | [Ibuprofen] |
| P:E | -2.05304 | 16.50328 | -0.1244 | 0.901775 | 1 | 89.44292 | 89.44292 | 0.437869 | 0.51289 | day3 | [Ibuprofen] |
| C:E | -2.05304 | 16.50328 | -0.1244 | 0.901775 | 1 | 67.0014 | 67.0014 | 0.328006 | 0.570838 | day3 | [Ibuprofen] |
| A:E | -34.9016 | 16.50328 | -2.11483 | 0.042325 | 1 | 410.5594 | 410.5594 | 2.009898 | 0.16594 | day3 | [Ibuprofen] |
| P:C:A | 22.6803 | 23.33917 | 0.97177 | 0.338454 | 1 | 333.5523 | 333.5523 | 1.632909 | 0.210494 | day3 | [Ibuprofen] |
| P:C:E | 2.053038 | 23.33917 | 0.087965 | 0.930452 | 1 | 1305.171 | 1305.171 | 6.389477 | 0.016615 | day3 | [Ibuprofen] |
| P:A:E | 34.90162 | 23.33917 | 1.49541 | 0.144605 | 1 | 58.97385 | 58.97385 | 0.288707 | 0.594767 | day3 | [Ibuprofen] |
| C:A:E | 55.27384 | 23.33917 | 2.368287 | 0.024078 | 1 | 99.26977 | 99.26977 | 0.485976 | 0.490762 | day3 | [Ibuprofen] |
| P:C:A:E | -87.5381 | 33.00657 | -2.65214 | 0.01234 | 1 | 1436.799 | 1436.799 | 7.033864 | 0.01234 | day3 | [Ibuprofen] |
| P | 6.531048 | 4.201948 | 1.554291 | 0.129951 | 1 | 1.965033 | 1.965033 | 0.074195 | 0.787073 | day4 | [Ibuprofen] |
| C | -2.03424 | 4.201948 | -0.48412 | 0.631598 | 1 | 851.696 | 851.696 | 32.15822 | 2.82E-06 | day4 | [Ibuprofen] |
| A | -8.10456 | 4.201948 | -1.92876 | 0.062671 | 1 | 848.9413 | 848.9413 | 32.05421 | 2.9E-06 | day4 | [Ibuprofen] |
| E | 3.845988 | 4.201948 | 0.915287 | 0.366887 | 1 | 18.48428 | 18.48428 | 0.697927 | 0.409676 | day4 | [Ibuprofen] |
| P:C | -2.56674 | 5.942451 | -0.43193 | 0.668687 | 1 | 342.4031 | 342.4031 | 12.92841 | 0.001074 | day4 | [Ibuprofen] |
| P:A | 7.554789 | 5.942451 | 1.271325 | 0.212776 | 1 | 7.976648 | 7.976648 | 0.301181 | 0.586952 | day4 | [Ibuprofen] |
| C:A | -9.51919 | 5.942451 | -1.6019 | 0.119006 | 1 | 443.4156 | 443.4156 | 16.74243 | 0.00027 | day4 | [Ibuprofen] |
| P:E | -7.02352 | 5.942451 | -1.18192 | 0.245945 | 1 | 1051.54 | 1051.54 | 39.7039 | 4.55E-07 | day4 | [Ibuprofen] |
| C:E | 2.975347 | 5.942451 | 0.500694 | 0.620013 | 1 | 14.20565 | 14.20565 | 0.536375 | 0.46927 | day4 | [Ibuprofen] |
| A:E | 6.090492 | 5.942451 | 1.024912 | 0.31309 | 1 | 0.050448 | 0.050448 | 0.001905 | 0.965459 | day4 | [Ibuprofen] |
| P:C:A | 2.962359 | 8.403895 | 0.352498 | 0.726776 | 1 | 150.5803 | 150.5803 | 5.685592 | 0.023202 | day4 | [Ibuprofen] |
| P:C:E | -2.0638 | 8.403895 | -0.24558 | 0.807578 | 1 | 276.354 | 276.354 | 10.43454 | 0.00286 | day4 | [Ibuprofen] |
| P:A:E | -4.20133 | 8.403895 | -0.49993 | 0.620547 | 1 | 341.3276 | 341.3276 | 12.8878 | 0.001091 | day4 | [Ibuprofen] |
| C:A:E | 8.892818 | 8.403895 | 1.058178 | 0.297897 | 1 | 50.91093 | 50.91093 | 1.922288 | 0.175189 | day4 | [Ibuprofen] |
| P:C:A:E | -34.2636 | 11.8849 | -2.88296 | 0.006986 | 1 | 220.1246 | 220.1246 | 8.311434 | 0.006986 | day4 | [Ibuprofen] |
| P | -0.84018 | 11.36446 | -0.07393 | 0.941526 | 1 | 165.0682 | 165.0682 | 0.852068 | 0.362876 | day7 | [Ibuprofen] |
| C | -0.38982 | 11.36446 | -0.0343 | 0.97285 | 1 | 133.2861 | 133.2861 | 0.688012 | 0.412986 | day7 | [Ibuprofen] |
| A | -5.90843 | 11.36446 | -0.5199 | 0.606709 | 1 | 156.1933 | 156.1933 | 0.806256 | 0.375936 | day7 | [Ibuprofen] |
| E | 0.01901 | 11.36446 | 0.001673 | 0.998676 | 1 | 284.3364 | 284.3364 | 1.46772 | 0.234576 | day7 | [Ibuprofen] |
| P:C | -2.44596 | 16.07178 | -0.15219 | 0.879993 | 1 | 149.9348 | 149.9348 | 0.773951 | 0.385557 | day7 | [Ibuprofen] |
| P:A | 7.384287 | 16.07178 | 0.459457 | 0.649012 | 1 | 402.9753 | 402.9753 | 2.080124 | 0.158945 | day7 | [Ibuprofen] |
| C:A | -24.2688 | 16.07178 | -1.51003 | 0.140849 | 1 | 1402.633 | 1402.633 | 7.240272 | 0.011237 | day7 | [Ibuprofen] |
| P:E | -34.9381 | 16.07178 | -2.17388 | 0.037222 | 1 | 1902.063 | 1902.063 | 9.818289 | 0.003684 | day7 | [Ibuprofen] |
| C:E | -1.8739 | 16.07178 | -0.1166 | 0.907909 | 1 | 120.0096 | 120.0096 | 0.619479 | 0.437032 | day7 | [Ibuprofen] |
| A:E | 4.092914 | 16.07178 | 0.254665 | 0.800612 | 1 | 146.233 | 146.233 | 0.754843 | 0.391417 | day7 | [Ibuprofen] |
| P:C:A | 29.00816 | 22.72893 | 1.276266 | 0.211047 | 1 | 333.3561 | 333.3561 | 1.720756 | 0.198929 | day7 | [Ibuprofen] |
| P:C:E | 40.11355 | 22.72893 | 1.764868 | 0.087129 | 1 | 74.65814 | 74.65814 | 0.385379 | 0.539135 | day7 | [Ibuprofen] |
| P:A:E | 29.49374 | 22.72893 | 1.29763 | 0.203691 | 1 | 318.1773 | 318.1773 | 1.642404 | 0.209205 | day7 | [Ibuprofen] |
| C:A:E | 26.3746 | 22.72893 | 1.160398 | 0.254471 | 1 | 421.8412 | 421.8412 | 2.177509 | 0.149814 | day7 | [Ibuprofen] |
| P:C:A:E | -100.181 | 32.14356 | -3.11669 | 0.003847 | 1 | 1881.811 | 1881.811 | 9.713751 | 0.003847 | day7 | [Ibuprofen] |
| P | -5.5E-14 | 5.43032 | -1E-14 | 1 | 1 | 435.25 | 435.25 | 9.840037 | 0.003651 | day11 | [Ibuprofen] |
| C | -7.6E-14 | 5.43032 | -1.4E-14 | 1 | 1 | 1421.412 | 1421.412 | 32.13496 | 2.84E-06 | day11 | [Ibuprofen] |
| A | -7.4E-14 | 5.43032 | -1.4E-14 | 1 | 1 | 2870.009 | 2870.009 | 64.88453 | 3.38E-09 | day11 | [Ibuprofen] |
| E | -7.1E-14 | 5.43032 | -1.3E-14 | 1 | 1 | 400.3739 | 400.3739 | 9.051567 | 0.005082 | day11 | [Ibuprofen] |
| P:C | -0.9334 | 7.679632 | -0.12154 | 0.904021 | 1 | 28.51248 | 28.51248 | 0.644604 | 0.427972 | day11 | [Ibuprofen] |
| P:A | -12.0049 | 7.679632 | -1.56321 | 0.127839 | 1 | 359.7313 | 359.7313 | 8.132726 | 0.007553 | day11 | [Ibuprofen] |
| C:A | -27.3318 | 7.679632 | -3.559 | 0.001187 | 1 | 452.3138 | 452.3138 | 10.22581 | 0.003115 | day11 | [Ibuprofen] |
| P:E | 5.91E-14 | 7.679632 | 7.7E-15 | 1 | 1 | 331.4462 | 331.4462 | 7.493264 | 0.010027 | day11 | [Ibuprofen] |
| C:E | -8.39345 | 7.679632 | -1.09295 | 0.282576 | 1 | 72.21031 | 72.21031 | 1.632515 | 0.210547 | day11 | [Ibuprofen] |
| A:E | -9.68901 | 7.679632 | -1.26165 | 0.216194 | 1 | 26.95797 | 26.95797 | 0.60946 | 0.440728 | day11 | [Ibuprofen] |
| P:C:A | 22.8086 | 10.86064 | 2.100116 | 0.04369 | 1 | 11.85869 | 11.85869 | 0.268099 | 0.608171 | day11 | [Ibuprofen] |
| P:C:E | -0.32258 | 10.86064 | -0.0297 | 0.976489 | 1 | 551.1148 | 551.1148 | 12.45948 | 0.001285 | day11 | [Ibuprofen] |
| P:A:E | 6.085468 | 10.86064 | 0.560323 | 0.579161 | 1 | 321.3523 | 321.3523 | 7.265063 | 0.011111 | day11 | [Ibuprofen] |
| C:A:E | 34.0822 | 10.86064 | 3.138139 | 0.003639 | 1 | 39.93703 | 39.93703 | 0.902888 | 0.34913 | day11 | [Ibuprofen] |
| P:C:A:E | -53.57 | 15.35926 | -3.48779 | 0.001439 | 1 | 538.0763 | 538.0763 | 12.16471 | 0.001439 | day11 | [Ibuprofen] |
| I | 10.37274 | 10.98252 | 0.944477 | 0.352003 | 1 | 1823.165 | 1823.165 | 10.07699 | 0.003311 | day3 | [Paracetamol] |
| C | 10.74635 | 10.98252 | 0.978496 | 0.335169 | 1 | 483.4716 | 483.4716 | 2.672243 | 0.111913 | day3 | [Paracetamol] |
| A | 5.20218 | 10.98252 | 0.473678 | 0.638945 | 1 | 927.2871 | 927.2871 | 5.125299 | 0.030494 | day3 | [Paracetamol] |
| E | 11.35827 | 10.98252 | 1.034214 | 0.308789 | 1 | 894.0133 | 894.0133 | 4.941388 | 0.033408 | day3 | [Paracetamol] |
| I:C | -5.13398 | 15.53162 | -0.33055 | 0.743139 | 1 | 116.2449 | 116.2449 | 0.642509 | 0.428716 | day3 | [Paracetamol] |
| I:A | -6.3811 | 15.53162 | -0.41085 | 0.683924 | 1 | 597.5035 | 597.5035 | 3.30252 | 0.078549 | day3 | [Paracetamol] |
| C:A | 7.507707 | 15.53162 | 0.483382 | 0.632116 | 1 | 13.03196 | 13.03196 | 0.07203 | 0.790126 | day3 | [Paracetamol] |
| I:E | 24.3675 | 15.53162 | 1.568896 | 0.126509 | 1 | 1255.912 | 1255.912 | 6.941673 | 0.012871 | day3 | [Paracetamol] |
| C:E | -4.53386 | 15.53162 | -0.29191 | 0.772238 | 1 | 112.8278 | 112.8278 | 0.623621 | 0.435518 | day3 | [Paracetamol] |
| A:E | -15.8335 | 15.53162 | -1.01944 | 0.31564 | 1 | 1738.538 | 1738.538 | 9.609246 | 0.004019 | day3 | [Paracetamol] |
| I:C:A | -10.078 | 21.96503 | -0.45882 | 0.649464 | 1 | 0.045734 | 0.045734 | 0.000253 | 0.987414 | day3 | [Paracetamol] |
| I:C:E | -2.4286 | 21.96503 | -0.11057 | 0.91265 | 1 | 46.76429 | 46.76429 | 0.258475 | 0.614657 | day3 | [Paracetamol] |
| I:A:E | -15.7101 | 21.96503 | -0.71523 | 0.479652 | 1 | 21.74982 | 21.74982 | 0.120216 | 0.731071 | day3 | [Paracetamol] |
| C:A:E | -11.0939 | 21.96503 | -0.50507 | 0.61697 | 1 | 0.443463 | 0.443463 | 0.002451 | 0.960822 | day3 | [Paracetamol] |
| I:C:A:E | 20.64991 | 31.06325 | 0.66477 | 0.51096 | 1 | 79.95351 | 79.95351 | 0.441919 | 0.51096 | day3 | [Paracetamol] |
| I | 9.272701 | 6.03842 | 1.535617 | 0.134462 | 1 | 309.0362 | 309.0362 | 5.650299 | 0.0236 | day4 | [Paracetamol] |
| C | 9.171779 | 6.03842 | 1.518904 | 0.138606 | 1 | 1487.486 | 1487.486 | 27.19661 | 1.06E-05 | day4 | [Paracetamol] |
| A | 3.091755 | 6.03842 | 0.512014 | 0.612157 | 1 | 28.39246 | 28.39246 | 0.519117 | 0.476449 | day4 | [Paracetamol] |
| E | 6.105431 | 6.03842 | 1.011098 | 0.319554 | 1 | 118.6651 | 118.6651 | 2.169626 | 0.150529 | day4 | [Paracetamol] |
| I:C | -8.10271 | 8.539615 | -0.94884 | 0.349814 | 1 | 139.6086 | 139.6086 | 2.55255 | 0.119946 | day4 | [Paracetamol] |
| I:A | -9.31608 | 8.539615 | -1.09092 | 0.283452 | 1 | 527.3014 | 527.3014 | 9.640976 | 0.003966 | day4 | [Paracetamol] |
| C:A | 37.23279 | 8.539615 | 4.360008 | 0.000126 | 1 | 678.1717 | 678.1717 | 12.39943 | 0.001315 | day4 | [Paracetamol] |
| I:E | 8.532315 | 8.539615 | 0.999145 | 0.32522 | 1 | 770.2183 | 770.2183 | 14.08238 | 0.000698 | day4 | [Paracetamol] |
| C:E | -7.44104 | 8.539615 | -0.87136 | 0.390051 | 1 | 994.3984 | 994.3984 | 18.1812 | 0.000166 | day4 | [Paracetamol] |
| A:E | -8.03584 | 8.539615 | -0.94101 | 0.353751 | 1 | 1731.403 | 1731.403 | 31.6563 | 3.21E-06 | day4 | [Paracetamol] |
| I:C:A | -14.0004 | 12.07684 | -1.15928 | 0.25492 | 1 | 29.7908 | 29.7908 | 0.544684 | 0.465876 | day4 | [Paracetamol] |
| I:C:E | 8.8644 | 12.07684 | 0.734 | 0.468293 | 1 | 205.734 | 205.734 | 3.761562 | 0.061302 | day4 | [Paracetamol] |
| I:A:E | -1.58082 | 12.07684 | -0.1309 | 0.896677 | 1 | 28.06466 | 28.06466 | 0.513123 | 0.478985 | day4 | [Paracetamol] |
| C:A:E | -38.0927 | 12.07684 | -3.1542 | 0.00349 | 1 | 692.8804 | 692.8804 | 12.66836 | 0.001186 | day4 | [Paracetamol] |
| I:C:A:E | 15.39593 | 17.07923 | 0.901442 | 0.374088 | 1 | 44.44402 | 44.44402 | 0.812597 | 0.374088 | day4 | [Paracetamol] |
| I | 2.52479 | 1.47449 | 1.712314 | 0.096515 | 1 | 2.791075 | 2.791075 | 0.855848 | 0.361828 | day7 | [Paracetamol] |
| C | 2.916888 | 1.47449 | 1.978235 | 0.056566 | 1 | 17.66697 | 17.66697 | 5.417356 | 0.026422 | day7 | [Paracetamol] |
| A | 1.013426 | 1.47449 | 0.687306 | 0.496841 | 1 | 1.62242 | 1.62242 | 0.497495 | 0.485706 | day7 | [Paracetamol] |
| E | -0.27923 | 1.47449 | -0.18937 | 0.850997 | 1 | 22.96334 | 22.96334 | 7.041421 | 0.012298 | day7 | [Paracetamol] |
| I:C | -3.41682 | 2.085243 | -1.63857 | 0.1111 | 1 | 5.735978 | 5.735978 | 1.758866 | 0.194155 | day7 | [Paracetamol] |
| I:A | -2.969 | 2.085243 | -1.42381 | 0.164179 | 1 | 5.384222 | 5.384222 | 1.651004 | 0.208045 | day7 | [Paracetamol] |
| C:A | 0.676529 | 2.085243 | 0.324437 | 0.747719 | 1 | 1.024484 | 1.024484 | 0.314145 | 0.579051 | day7 | [Paracetamol] |
| I:E | -2.52479 | 2.085243 | -1.21079 | 0.234843 | 1 | 0.205712 | 0.205712 | 0.063079 | 0.8033 | day7 | [Paracetamol] |
| C:E | -1.84177 | 2.085243 | -0.88324 | 0.383695 | 1 | 1.80927 | 1.80927 | 0.55479 | 0.461803 | day7 | [Paracetamol] |
| A:E | 0.230761 | 2.085243 | 0.110664 | 0.912574 | 1 | 2.382647 | 2.382647 | 0.730609 | 0.39904 | day7 | [Paracetamol] |
| I:C:A | 0.219651 | 2.94898 | 0.074484 | 0.941089 | 1 | 1.764468 | 1.764468 | 0.541052 | 0.467355 | day7 | [Paracetamol] |
| I:C:E | 2.534313 | 2.94898 | 0.859386 | 0.39652 | 1 | 11.10815 | 11.10815 | 3.406174 | 0.074223 | day7 | [Paracetamol] |
| I:A:E | 1.724811 | 2.94898 | 0.584884 | 0.562727 | 1 | 6.926581 | 6.926581 | 2.123949 | 0.154757 | day7 | [Paracetamol] |
| C:A:E | -1.71813 | 2.94898 | -0.58262 | 0.564232 | 1 | 0.122387 | 0.122387 | 0.037528 | 0.847618 | day7 | [Paracetamol] |
| I:C:A:E | 2.628351 | 4.170487 | 0.630227 | 0.533021 | 1 | 1.295293 | 1.295293 | 0.397185 | 0.533021 | day7 | [Paracetamol] |
| I | 1.31E-15 | 0.814423 | 1.61E-15 | 1 | 1 | 0.744753 | 0.744753 | 0.74855 | 0.393376 | day11 | [Paracetamol] |
| C | 2.972897 | 0.814423 | 3.65031 | 0.000925 | 1 | 4.821516 | 4.821516 | 4.846099 | 0.035038 | day11 | [Paracetamol] |
| A | -1E-16 | 0.814423 | -1.3E-16 | 1 | 1 | 3.176375 | 3.176375 | 3.19257 | 0.083451 | day11 | [Paracetamol] |
| E | -3.9E-16 | 0.814423 | -4.8E-16 | 1 | 1 | 4.516345 | 4.516345 | 4.539372 | 0.040907 | day11 | [Paracetamol] |
| I:C | -1.43391 | 1.151768 | -1.24496 | 0.222186 | 1 | 0.928392 | 0.928392 | 0.933126 | 0.341296 | day11 | [Paracetamol] |
| I:A | 0.116094 | 1.151768 | 0.100796 | 0.920341 | 1 | 0.302472 | 0.302472 | 0.304015 | 0.585206 | day11 | [Paracetamol] |
| C:A | -2.49536 | 1.151768 | -2.16655 | 0.037824 | 1 | 3.544857 | 3.544857 | 3.562931 | 0.068181 | day11 | [Paracetamol] |
| I:E | -8.5E-16 | 1.151768 | -7.4E-16 | 1 | 1 | 0.478622 | 0.478622 | 0.481062 | 0.492946 | day11 | [Paracetamol] |
| C:E | -2.77525 | 1.151768 | -2.40956 | 0.021898 | 1 | 4.099123 | 4.099123 | 4.120023 | 0.050762 | day11 | [Paracetamol] |
| A:E | -2.5E-15 | 1.151768 | -2.2E-15 | 1 | 1 | 2.595556 | 2.595556 | 2.60879 | 0.116091 | day11 | [Paracetamol] |
| I:C:A | 0.84028 | 1.628846 | 0.515874 | 0.609489 | 1 | 0.201992 | 0.201992 | 0.203021 | 0.655331 | day11 | [Paracetamol] |
| I:C:E | 1.236262 | 1.628846 | 0.75898 | 0.453419 | 1 | 0.627843 | 0.627843 | 0.631044 | 0.432825 | day11 | [Paracetamol] |
| I:A:E | -0.11609 | 1.628846 | -0.07127 | 0.943624 | 1 | 0.143496 | 0.143496 | 0.144228 | 0.706622 | day11 | [Paracetamol] |
| C:A:E | 2.297718 | 1.628846 | 1.410642 | 0.167998 | 1 | 2.92962 | 2.92962 | 2.944557 | 0.095835 | day11 | [Paracetamol] |
| I:C:A:E | -0.64263 | 2.303536 | -0.27898 | 0.782057 | 1 | 0.077434 | 0.077434 | 0.077828 | 0.782057 | day11 | [Paracetamol] |
| I | -18.609 | 20.06217 | -0.92757 | 0.360799 | 1 | 0.427638 | 0.427638 | 0.000708 | 0.978938 | day3 | [Caffein] |
| P | 41.46054 | 20.06217 | 2.066603 | 0.047211 | 1 | 34195.96 | 34195.96 | 56.64059 | 1.76E-08 | day3 | [Caffein] |
| A | -18.2601 | 20.06217 | -0.91017 | 0.369754 | 1 | 867.4753 | 867.4753 | 1.436845 | 0.239734 | day3 | [Caffein] |
| E | -34.8996 | 20.06217 | -1.73957 | 0.091856 | 1 | 867.9761 | 867.9761 | 1.437675 | 0.239601 | day3 | [Caffein] |
| I:P | 20.16121 | 28.37219 | 0.710598 | 0.482646 | 1 | 283.664 | 283.664 | 0.469848 | 0.498151 | day3 | [Caffein] |
| I:A | -12.2909 | 28.37219 | -0.4332 | 0.667868 | 1 | 456.0944 | 456.0944 | 0.755453 | 0.391435 | day3 | [Caffein] |
| P:A | 19.00177 | 28.37219 | 0.669732 | 0.507987 | 1 | 684.4787 | 684.4787 | 1.133738 | 0.295203 | day3 | [Caffein] |
| I:E | 70.10542 | 28.37219 | 2.47092 | 0.019177 | 1 | 6625.36 | 6625.36 | 10.97394 | 0.002357 | day3 | [Caffein] |
| P:E | 27.78317 | 28.37219 | 0.979239 | 0.335042 | 1 | 5.839216 | 5.839216 | 0.009672 | 0.922291 | day3 | [Caffein] |
| A:E | 20.22474 | 30.09326 | 0.672069 | 0.506518 | 1 | 2004.119 | 2004.119 | 3.319529 | 0.078123 | day3 | [Caffein] |
| I:P:A | -28.0627 | 40.12434 | -0.69939 | 0.489521 | 1 | 164.5456 | 164.5456 | 0.272546 | 0.60534 | day3 | [Caffein] |
| I:P:E | -75.8361 | 40.12434 | -1.89003 | 0.068136 | 1 | 827.7633 | 827.7633 | 1.371068 | 0.250551 | day3 | [Caffein] |
| I:A:E | -11.4744 | 41.35922 | -0.27743 | 0.783289 | 1 | 696.4714 | 696.4714 | 1.153603 | 0.291086 | day3 | [Caffein] |
| P:A:E | -19.2614 | 41.35922 | -0.46571 | 0.644679 | 1 | 413.4553 | 413.4553 | 0.684828 | 0.414255 | day3 | [Caffein] |
| I:P:A:E | 83.65244 | 57.6242 | 1.45169 | 0.156634 | 1 | 1272.315 | 1272.315 | 2.107402 | 0.156634 | day3 | [Caffein] |
| I | 3.19E-14 | 9.394534 | 3.39E-15 | 1 | 1 | 905.4607 | 905.4607 | 6.839556 | 0.013647 | day4 | [Caffein] |
| P | 59.12361 | 9.394534 | 6.293406 | 5.34E-07 | 1 | 749.2719 | 749.2719 | 5.659756 | 0.0237 | day4 | [Caffein] |
| A | -7.6E-15 | 9.394534 | -8.1E-16 | 1 | 1 | 119.1103 | 119.1103 | 0.89972 | 0.350193 | day4 | [Caffein] |
| E | 1.39E-14 | 9.394534 | 1.48E-15 | 1 | 1 | 1518.041 | 1518.041 | 11.46679 | 0.001941 | day4 | [Caffein] |
| I:P | -59.1236 | 13.28588 | -4.45011 | 0.000103 | 1 | 2379.002 | 2379.002 | 17.97021 | 0.000187 | day4 | [Caffein] |
| I:A | 12.23541 | 13.28588 | 0.920934 | 0.364197 | 1 | 226.9577 | 226.9577 | 1.714364 | 0.200042 | day4 | [Caffein] |
| P:A | -29.4761 | 13.28588 | -2.2186 | 0.03397 | 1 | 149.6595 | 149.6595 | 1.130479 | 0.295886 | day4 | [Caffein] |
| I:E | 10.15218 | 13.28588 | 0.764133 | 0.450565 | 1 | 1545.397 | 1545.397 | 11.67343 | 0.00179 | day4 | [Caffein] |
| P:E | -59.1236 | 13.28588 | -4.45011 | 0.000103 | 1 | 1276.101 | 1276.101 | 9.639251 | 0.004048 | day4 | [Caffein] |
| A:E | -1.8E-14 | 14.0918 | -1.3E-15 | 1 | 1 | 3.045312 | 3.045312 | 0.023003 | 0.880431 | day4 | [Caffein] |
| I:P:A | 17.24066 | 18.78907 | 0.91759 | 0.365918 | 1 | 114.6231 | 114.6231 | 0.865826 | 0.359304 | day4 | [Caffein] |
| I:P:E | 48.97144 | 18.78907 | 2.606379 | 0.01394 | 1 | 1511.446 | 1511.446 | 11.41697 | 0.001979 | day4 | [Caffein] |
| I:A:E | -22.3876 | 19.36733 | -1.15595 | 0.256535 | 1 | 523.2411 | 523.2411 | 3.952393 | 0.055697 | day4 | [Caffein] |
| P:A:E | 29.47607 | 19.36733 | 1.521948 | 0.138158 | 1 | 485.4647 | 485.4647 | 3.667042 | 0.064765 | day4 | [Caffein] |
| I:P:A:E | -7.08848 | 26.98374 | -0.26269 | 0.794524 | 1 | 9.135742 | 9.135742 | 0.069008 | 0.794524 | day4 | [Caffein] |
| I | 0 | 0 |  |  | 1 | 0 | 0 |  |  | day7 | [Caffein] |
| P | 0 | 0 |  |  | 1 | 0 | 0 |  |  | day7 | [Caffein] |
| A | 0 | 0 |  |  | 1 | 0 | 0 |  |  | day7 | [Caffein] |
| E | 0 | 0 |  |  | 1 | 0 | 0 |  |  | day7 | [Caffein] |
| I:P | 0 | 0 |  |  | 1 | 0 | 0 |  |  | day7 | [Caffein] |
| I:A | 0 | 0 |  |  | 1 | 0 | 0 |  |  | day7 | [Caffein] |
| P:A | 0 | 0 |  |  | 1 | 0 | 0 |  |  | day7 | [Caffein] |
| I:E | 0 | 0 |  |  | 1 | 0 | 0 |  |  | day7 | [Caffein] |
| P:E | 0 | 0 |  |  | 1 | 0 | 0 |  |  | day7 | [Caffein] |
| A:E | 0 | 0 |  |  | 1 | 0 | 0 |  |  | day7 | [Caffein] |
| I:P:A | 0 | 0 |  |  | 1 | 0 | 0 |  |  | day7 | [Caffein] |
| I:P:E | 0 | 0 |  |  | 1 | 0 | 0 |  |  | day7 | [Caffein] |
| I:A:E | 0 | 0 |  |  | 1 | 0 | 0 |  |  | day7 | [Caffein] |
| P:A:E | 0 | 0 |  |  | 1 | 0 | 0 |  |  | day7 | [Caffein] |
| I:P:A:E | 0 | 0 |  |  | 1 | 0 | 0 |  |  | day7 | [Caffein] |
| I | 0 | 0 |  |  | 1 | 0 | 0 |  |  | day11 | [Caffein] |
| P | 0 | 0 |  |  | 1 | 0 | 0 |  |  | day11 | [Caffein] |
| A | 0 | 0 |  |  | 1 | 0 | 0 |  |  | day11 | [Caffein] |
| E | 0 | 0 |  |  | 1 | 0 | 0 |  |  | day11 | [Caffein] |
| I:P | 0 | 0 |  |  | 1 | 0 | 0 |  |  | day11 | [Caffein] |
| I:A | 0 | 0 |  |  | 1 | 0 | 0 |  |  | day11 | [Caffein] |
| P:A | 0 | 0 |  |  | 1 | 0 | 0 |  |  | day11 | [Caffein] |
| I:E | 0 | 0 |  |  | 1 | 0 | 0 |  |  | day11 | [Caffein] |
| P:E | 0 | 0 |  |  | 1 | 0 | 0 |  |  | day11 | [Caffein] |
| A:E | 0 | 0 |  |  | 1 | 0 | 0 |  |  | day11 | [Caffein] |
| I:P:A | 0 | 0 |  |  | 1 | 0 | 0 |  |  | day11 | [Caffein] |
| I:P:E | 0 | 0 |  |  | 1 | 0 | 0 |  |  | day11 | [Caffein] |
| I:A:E | 0 | 0 |  |  | 1 | 0 | 0 |  |  | day11 | [Caffein] |
| P:A:E | 0 | 0 |  |  | 1 | 0 | 0 |  |  | day11 | [Caffein] |
| I:P:A:E | 0 | 0 |  |  | 1 | 0 | 0 |  |  | day11 | [Caffein] |
| I | -3.46376 | 2.90291 | -1.1932 | 0.241842 | 1 | 53.71593 | 53.71593 | 4.249568 | 0.047731 | day3 | [Enalapril] |
| P | -2.30902 | 2.90291 | -0.79542 | 0.43242 | 1 | 213.7546 | 213.7546 | 16.91053 | 0.000267 | day3 | [Enalapril] |
| A | -0.40292 | 2.90291 | -0.1388 | 0.890507 | 1 | 8.621164 | 8.621164 | 0.682037 | 0.415197 | day3 | [Enalapril] |
| C | -2.49139 | 2.90291 | -0.85824 | 0.397348 | 1 | 20.78976 | 20.78976 | 1.644717 | 0.209187 | day3 | [Enalapril] |
| I:P | 5.008145 | 4.105334 | 1.219912 | 0.231698 | 1 | 0.445153 | 0.445153 | 0.035217 | 0.852365 | day3 | [Enalapril] |
| I:A | 4.797251 | 4.105334 | 1.168541 | 0.251497 | 1 | 79.95534 | 79.95534 | 6.325417 | 0.0173 | day3 | [Enalapril] |
| P:A | 1.347786 | 4.105334 | 0.328301 | 0.744891 | 1 | 35.26707 | 35.26707 | 2.790044 | 0.10492 | day3 | [Enalapril] |
| I:C | 0.633089 | 4.105334 | 0.154211 | 0.878443 | 1 | 16.05156 | 16.05156 | 1.269869 | 0.268439 | day3 | [Enalapril] |
| P:C | 4.233729 | 4.105334 | 1.031275 | 0.310391 | 1 | 175.7013 | 175.7013 | 13.90006 | 0.000773 | day3 | [Enalapril] |
| A:C | -17.0814 | 4.354365 | -3.92283 | 0.000452 | 1 | 42.23163 | 42.23163 | 3.341024 | 0.077212 | day3 | [Enalapril] |
| I:P:A | -0.91718 | 5.805819 | -0.15798 | 0.875501 | 1 | 136.1157 | 136.1157 | 10.76837 | 0.002558 | day3 | [Enalapril] |
| I:P:C | 1.359633 | 5.805819 | 0.234185 | 0.816382 | 1 | 99.91659 | 99.91659 | 7.904589 | 0.008474 | day3 | [Enalapril] |
| I:A:C | 17.85379 | 5.984502 | 2.983338 | 0.005517 | 1 | 8.671462 | 8.671462 | 0.686016 | 0.413854 | day3 | [Enalapril] |
| P:A:C | 21.07566 | 5.984502 | 3.521708 | 0.001352 | 1 | 34.39635 | 34.39635 | 2.72116 | 0.109124 | day3 | [Enalapril] |
| I:P:A:C | -27.568 | 8.337973 | -3.30632 | 0.002397 | 1 | 138.1807 | 138.1807 | 10.93173 | 0.002397 | day3 | [Enalapril] |
| I | -6.27102 | 4.096895 | -1.53068 | 0.135991 | 1 | 17.38556 | 17.38556 | 0.690538 | 0.412336 | day4 | [Enalapril] |
| P | -7.03855 | 4.096895 | -1.71802 | 0.095767 | 1 | 394.3865 | 394.3865 | 15.66467 | 0.000411 | day4 | [Enalapril] |
| A | -6.45773 | 4.096895 | -1.57625 | 0.12512 | 1 | 167.7214 | 167.7214 | 6.661741 | 0.014806 | day4 | [Enalapril] |
| C | -20.158 | 4.096895 | -4.92031 | 2.7E-05 | 1 | 265.4242 | 265.4242 | 10.54241 | 0.002801 | day4 | [Enalapril] |
| I:P | 6.846896 | 5.793884 | 1.181745 | 0.246293 | 1 | 18.00076 | 18.00076 | 0.714974 | 0.404277 | day4 | [Enalapril] |
| I:A | 8.345816 | 5.793884 | 1.440453 | 0.159764 | 1 | 123.6629 | 123.6629 | 4.911776 | 0.034146 | day4 | [Enalapril] |
| P:A | 8.707478 | 5.793884 | 1.502874 | 0.142991 | 1 | 1.076931 | 1.076931 | 0.042775 | 0.837503 | day4 | [Enalapril] |
| I:C | 6.600356 | 5.793884 | 1.139194 | 0.263349 | 1 | 24.32961 | 24.32961 | 0.96635 | 0.333203 | day4 | [Enalapril] |
| P:C | 22.94884 | 5.793884 | 3.960874 | 0.000407 | 1 | 517.5044 | 517.5044 | 20.5548 | 8.13E-05 | day4 | [Enalapril] |
| A:C | 5.394337 | 6.145342 | 0.877793 | 0.386809 | 1 | 79.16772 | 79.16772 | 3.144469 | 0.086011 | day4 | [Enalapril] |
| I:P:A | -5.00383 | 8.193789 | -0.61069 | 0.545859 | 1 | 87.06056 | 87.06056 | 3.457965 | 0.072461 | day4 | [Enalapril] |
| I:P:C | -8.91494 | 8.193789 | -1.08801 | 0.284977 | 1 | 169.7465 | 169.7465 | 6.742173 | 0.014269 | day4 | [Enalapril] |
| I:A:C | 8.891539 | 8.445965 | 1.052756 | 0.30059 | 1 | 5.258245 | 5.258245 | 0.208853 | 0.650855 | day4 | [Enalapril] |
| P:A:C | -3.64456 | 8.445965 | -0.43151 | 0.669081 | 1 | 74.966 | 74.966 | 2.977581 | 0.094382 | day4 | [Enalapril] |
| I:P:A:C | -12.6245 | 11.76743 | -1.07283 | 0.291627 | 1 | 28.97779 | 28.97779 | 1.150971 | 0.291627 | day4 | [Enalapril] |
| I | -2.50341 | 18.29563 | -0.13683 | 0.892049 | 1 | 611.8724 | 611.8724 | 1.218639 | 0.278123 | day7 | [Enalapril] |
| P | -4.55614 | 18.29563 | -0.24903 | 0.804981 | 1 | 408.6693 | 408.6693 | 0.813929 | 0.373918 | day7 | [Enalapril] |
| A | -7.41292 | 18.29563 | -0.40517 | 0.688132 | 1 | 489.5927 | 489.5927 | 0.9751 | 0.331057 | day7 | [Enalapril] |
| C | -5.15506 | 18.29563 | -0.28176 | 0.779996 | 1 | 190.6919 | 190.6919 | 0.379793 | 0.542214 | day7 | [Enalapril] |
| I:P | -49.3361 | 25.87392 | -1.90679 | 0.065853 | 1 | 168.2494 | 168.2494 | 0.335095 | 0.566857 | day7 | [Enalapril] |
| I:A | -1.52899 | 25.87392 | -0.05909 | 0.953257 | 1 | 2517.073 | 2517.073 | 5.013142 | 0.032472 | day7 | [Enalapril] |
| P:A | -23.5645 | 25.87392 | -0.91074 | 0.369458 | 1 | 90.9914 | 90.9914 | 0.181224 | 0.673265 | day7 | [Enalapril] |
| I:C | -33.5115 | 25.87392 | -1.29518 | 0.20482 | 1 | 251.8573 | 251.8573 | 0.501613 | 0.484084 | day7 | [Enalapril] |
| P:C | -2.63685 | 25.87392 | -0.10191 | 0.919483 | 1 | 4519.337 | 4519.337 | 9.000962 | 0.005287 | day7 | [Enalapril] |
| A:C | -27.1943 | 27.44344 | -0.99092 | 0.329396 | 1 | 214.3933 | 214.3933 | 0.426998 | 0.518284 | day7 | [Enalapril] |
| I:P:A | 66.52103 | 36.59125 | 1.817949 | 0.078748 | 1 | 294.7157 | 294.7157 | 0.586972 | 0.449387 | day7 | [Enalapril] |
| I:P:C | 89.67454 | 36.59125 | 2.45071 | 0.020098 | 1 | 1443.006 | 1443.006 | 2.873972 | 0.100046 | day7 | [Enalapril] |
| I:A:C | 43.95274 | 37.7174 | 1.165317 | 0.252779 | 1 | 8.949043 | 8.949043 | 0.017823 | 0.894657 | day7 | [Enalapril] |
| P:A:C | 40.2692 | 37.7174 | 1.067656 | 0.29392 | 1 | 39.92758 | 39.92758 | 0.079522 | 0.77982 | day7 | [Enalapril] |
| I:P:A:C | -92.5461 | 52.55018 | -1.7611 | 0.088084 | 1 | 1557.234 | 1557.234 | 3.101473 | 0.088084 | day7 | [Enalapril] |
| I | -5.83798 | 10.27133 | -0.56838 | 0.573874 | 1 | 497.5082 | 497.5082 | 3.143807 | 0.086042 | day11 | [Enalapril] |
| P | -83.3769 | 10.27133 | -8.11745 | 3.62E-09 | 1 | 15580.5 | 15580.5 | 98.45481 | 3.86E-11 | day11 | [Enalapril] |
| A | -25.2146 | 10.27133 | -2.45485 | 0.019906 | 1 | 2969.309 | 2969.309 | 18.76338 | 0.000144 | day11 | [Enalapril] |
| C | -22.6968 | 10.27133 | -2.20972 | 0.034641 | 1 | 4698.051 | 4698.051 | 29.68748 | 5.93E-06 | day11 | [Enalapril] |
| I:P | 20.01005 | 14.52585 | 1.377548 | 0.178208 | 1 | 1086.139 | 1086.139 | 6.863428 | 0.0135 | day11 | [Enalapril] |
| I:A | -12.2348 | 14.52585 | -0.84228 | 0.406084 | 1 | 23.18247 | 23.18247 | 0.146492 | 0.704522 | day11 | [Enalapril] |
| P:A | 37.8888 | 14.52585 | 2.60837 | 0.013873 | 1 | 4186.002 | 4186.002 | 26.45179 | 1.42E-05 | day11 | [Enalapril] |
| I:C | -27.0577 | 14.52585 | -1.86273 | 0.072001 | 1 | 125.7165 | 125.7165 | 0.794416 | 0.379634 | day11 | [Enalapril] |
| P:C | 30.04948 | 14.52585 | 2.06869 | 0.047002 | 1 | 4960.132 | 4960.132 | 31.34359 | 3.86E-06 | day11 | [Enalapril] |
| A:C | -37.179 | 15.40699 | -2.41313 | 0.021919 | 1 | 149.5529 | 149.5529 | 0.94504 | 0.33851 | day11 | [Enalapril] |
| I:P:A | -4.56744 | 20.54265 | -0.22234 | 0.825508 | 1 | 355.1072 | 355.1072 | 2.24396 | 0.144252 | day11 | [Enalapril] |
| I:P:C | 16.07666 | 20.54265 | 0.782599 | 0.4398 | 1 | 1.545784 | 1.545784 | 0.009768 | 0.921907 | day11 | [Enalapril] |
| I:A:C | 48.88596 | 21.17488 | 2.308677 | 0.027795 | 1 | 607.5753 | 607.5753 | 3.839332 | 0.059109 | day11 | [Enalapril] |
| P:A:C | 28.05745 | 21.17488 | 1.325035 | 0.194844 | 1 | 50.06572 | 50.06572 | 0.316371 | 0.577842 | day11 | [Enalapril] |
| I:P:A:C | -38.3659 | 29.50214 | -1.30044 | 0.203034 | 1 | 267.626 | 267.626 | 1.691157 | 0.203034 | day11 | [Enalapril] |
| I | 10.4025 | 6.722406 | 1.547438 | 0.131907 | 1 | 273.0671 | 273.0671 | 4.028364 | 0.053529 | day3 | [Atenolol] |
| P | 4.68147 | 6.722406 | 0.696398 | 0.491369 | 1 | 47.3009 | 47.3009 | 0.697796 | 0.409917 | day3 | [Atenolol] |
| E | 3.245899 | 6.722406 | 0.482848 | 0.632596 | 1 | 240.6171 | 240.6171 | 3.549651 | 0.068967 | day3 | [Atenolol] |
| C | 1.684829 | 6.722406 | 0.250629 | 0.803755 | 1 | 1.739519 | 1.739519 | 0.025662 | 0.873768 | day3 | [Atenolol] |
| I:P | -6.79329 | 9.506917 | -0.71456 | 0.480226 | 1 | 191.5582 | 191.5582 | 2.825921 | 0.102804 | day3 | [Atenolol] |
| I:E | 0.197685 | 9.506917 | 0.020794 | 0.983543 | 1 | 14.3174 | 14.3174 | 0.211214 | 0.649024 | day3 | [Atenolol] |
| P:E | -1.07334 | 9.506917 | -0.1129 | 0.910837 | 1 | 107.4437 | 107.4437 | 1.58504 | 0.217435 | day3 | [Atenolol] |
| I:C | -1.11616 | 9.506917 | -0.1174 | 0.907297 | 1 | 122.5744 | 122.5744 | 1.808253 | 0.188475 | day3 | [Atenolol] |
| P:C | 0.909554 | 9.506917 | 0.095673 | 0.924396 | 1 | 0.843104 | 0.843104 | 0.012438 | 0.911919 | day3 | [Atenolol] |
| E:C | -1.99602 | 10.08361 | -0.19795 | 0.844379 | 1 | 4.039533 | 4.039533 | 0.059592 | 0.80875 | day3 | [Atenolol] |
| I:P:E | 4.075809 | 13.44481 | 0.303151 | 0.7638 | 1 | 49.46319 | 49.46319 | 0.729695 | 0.399535 | day3 | [Atenolol] |
| I:P:C | -10.6375 | 13.44481 | -0.7912 | 0.434842 | 1 | 32.39513 | 32.39513 | 0.477902 | 0.494519 | day3 | [Atenolol] |
| I:E:C | -3.76641 | 13.85859 | -0.27177 | 0.787597 | 1 | 0.017107 | 0.017107 | 0.000252 | 0.987427 | day3 | [Atenolol] |
| P:E:C | 6.175671 | 13.85859 | 0.44562 | 0.658968 | 1 | 74.04397 | 74.04397 | 1.092318 | 0.30404 | day3 | [Atenolol] |
| I:P:E:C | 7.589635 | 19.30864 | 0.393069 | 0.696958 | 1 | 10.47319 | 10.47319 | 0.154504 | 0.696958 | day3 | [Atenolol] |
| I | 6.949261 | 7.06821 | 0.983171 | 0.333135 | 1 | 804.8784 | 804.8784 | 10.74039 | 0.002587 | day4 | [Atenolol] |
| P | 9.487114 | 7.06821 | 1.342223 | 0.189272 | 1 | 746.6294 | 746.6294 | 9.963109 | 0.003543 | day4 | [Atenolol] |
| E | 2.930448 | 7.06821 | 0.414595 | 0.681293 | 1 | 112.6057 | 112.6057 | 1.502624 | 0.229499 | day4 | [Atenolol] |
| C | 2.236841 | 7.06821 | 0.316465 | 0.753771 | 1 | 142.7286 | 142.7286 | 1.904587 | 0.177438 | day4 | [Atenolol] |
| I:P | -0.03921 | 9.995959 | -0.00392 | 0.996896 | 1 | 133.9152 | 133.9152 | 1.78698 | 0.191023 | day4 | [Atenolol] |
| I:E | 8.707527 | 9.995959 | 0.871105 | 0.390393 | 1 | 3.503748 | 3.503748 | 0.046754 | 0.830227 | day4 | [Atenolol] |
| P:E | 1.04777 | 9.995959 | 0.104819 | 0.917194 | 1 | 3.326576 | 3.326576 | 0.04439 | 0.834508 | day4 | [Atenolol] |
| I:C | 9.28882 | 9.995959 | 0.929258 | 0.359936 | 1 | 4.94731 | 4.94731 | 0.066017 | 0.798924 | day4 | [Atenolol] |
| P:C | 3.206088 | 9.995959 | 0.320738 | 0.750561 | 1 | 0.335683 | 0.335683 | 0.004479 | 0.947069 | day4 | [Atenolol] |
| E:C | 1.057544 | 10.60232 | 0.099746 | 0.921188 | 1 | 69.71291 | 69.71291 | 0.930257 | 0.34226 | day4 | [Atenolol] |
| I:P:E | -9.17288 | 14.13642 | -0.64888 | 0.521193 | 1 | 6.552628 | 6.552628 | 0.087439 | 0.76943 | day4 | [Atenolol] |
| I:P:C | -10.6319 | 14.13642 | -0.75209 | 0.457668 | 1 | 15.31472 | 15.31472 | 0.204361 | 0.654371 | day4 | [Atenolol] |
| I:E:C | -16.7849 | 14.57149 | -1.1519 | 0.258169 | 1 | 78.19756 | 78.19756 | 1.043477 | 0.31492 | day4 | [Atenolol] |
| P:E:C | -1.35798 | 14.57149 | -0.09319 | 0.926349 | 1 | 19.83854 | 19.83854 | 0.264728 | 0.61054 | day4 | [Atenolol] |
| I:P:E:C | 12.76987 | 20.30189 | 0.628999 | 0.533958 | 1 | 29.64901 | 29.64901 | 0.39564 | 0.533958 | day4 | [Atenolol] |
| I | 29.23774 | 6.73282 | 4.34257 | 0.00014 | 1 | 2539.652 | 2539.652 | 37.34986 | 8.94E-07 | day7 | [Atenolol] |
| P | 42.22152 | 6.73282 | 6.271001 | 5.69E-07 | 1 | 6689.1 | 6689.1 | 98.37449 | 3.9E-11 | day7 | [Atenolol] |
| E | 6.327245 | 6.73282 | 0.939761 | 0.354605 | 1 | 120.3208 | 120.3208 | 1.76952 | 0.193145 | day7 | [Atenolol] |
| C | 3.091931 | 6.73282 | 0.459233 | 0.649271 | 1 | 60.02175 | 60.02175 | 0.882721 | 0.354721 | day7 | [Atenolol] |
| I:P | -27.4165 | 9.521645 | -2.87939 | 0.007162 | 1 | 1793.548 | 1793.548 | 26.37714 | 1.45E-05 | day7 | [Atenolol] |
| I:E | 7.540421 | 9.521645 | 0.791924 | 0.434423 | 1 | 157.0196 | 157.0196 | 2.309238 | 0.138742 | day7 | [Atenolol] |
| P:E | -17.0112 | 9.521645 | -1.78659 | 0.083789 | 1 | 98.22393 | 98.22393 | 1.444548 | 0.238506 | day7 | [Atenolol] |
| I:C | -10.0759 | 9.521645 | -1.05821 | 0.298137 | 1 | 217.6329 | 217.6329 | 3.200659 | 0.083385 | day7 | [Atenolol] |
| P:C | -5.77581 | 9.521645 | -0.6066 | 0.548535 | 1 | 161.9184 | 161.9184 | 2.381282 | 0.132945 | day7 | [Atenolol] |
| E:C | -9.31 | 10.09923 | -0.92185 | 0.363725 | 1 | 8.976141 | 8.976141 | 0.132009 | 0.718824 | day7 | [Atenolol] |
| I:P:E | 1.655234 | 13.46564 | 0.122923 | 0.902962 | 1 | 0.059304 | 0.059304 | 0.000872 | 0.976629 | day7 | [Atenolol] |
| I:P:C | 5.70549 | 13.46564 | 0.423707 | 0.674705 | 1 | 10.93623 | 10.93623 | 0.160836 | 0.691139 | day7 | [Atenolol] |
| I:E:C | 0.451737 | 13.88006 | 0.032546 | 0.974245 | 1 | 5.282592 | 5.282592 | 0.077689 | 0.782304 | day7 | [Atenolol] |
| P:E:C | 23.60681 | 13.88006 | 1.700771 | 0.098998 | 1 | 323.5477 | 323.5477 | 4.758313 | 0.036864 | day7 | [Atenolol] |
| I:P:E:C | -4.90023 | 19.33855 | -0.25339 | 0.801638 | 1 | 4.365868 | 4.365868 | 0.064207 | 0.801638 | day7 | [Atenolol] |
| I | 61.50499 | 14.64725 | 4.199081 | 0.000209 | 1 | 10402.49 | 10402.49 | 32.32466 | 3.01E-06 | day11 | [Atenolol] |
| P | 30.54476 | 14.64725 | 2.085358 | 0.045362 | 1 | 2210.05 | 2210.05 | 6.867499 | 0.013475 | day11 | [Atenolol] |
| E | 6.694092 | 14.64725 | 0.45702 | 0.650843 | 1 | 26.01772 | 26.01772 | 0.080847 | 0.778043 | day11 | [Atenolol] |
| C | -9.49105 | 14.64725 | -0.64797 | 0.521772 | 1 | 1565.329 | 1565.329 | 4.864096 | 0.034966 | day11 | [Atenolol] |
| I:P | -45.4137 | 20.71434 | -2.19238 | 0.035985 | 1 | 3538.689 | 3538.689 | 10.9961 | 0.002336 | day11 | [Atenolol] |
| I:E | -5.357 | 20.71434 | -0.25861 | 0.797643 | 1 | 0.479509 | 0.479509 | 0.00149 | 0.969456 | day11 | [Atenolol] |
| P:E | -24.0259 | 20.71434 | -1.15987 | 0.254959 | 1 | 3.07701 | 3.07701 | 0.009561 | 0.922734 | day11 | [Atenolol] |
| I:C | -20.0856 | 20.71434 | -0.96965 | 0.339728 | 1 | 1414.663 | 1414.663 | 4.395917 | 0.04428 | day11 | [Atenolol] |
| P:C | 10.56304 | 20.71434 | 0.509939 | 0.613706 | 1 | 2068.671 | 2068.671 | 6.428176 | 0.016493 | day11 | [Atenolol] |
| E:C | -10.9687 | 21.97088 | -0.49924 | 0.621137 | 1 | 2.281741 | 2.281741 | 0.00709 | 0.933436 | day11 | [Atenolol] |
| I:P:E | 21.41128 | 29.2945 | 0.730898 | 0.47033 | 1 | 149.2052 | 149.2052 | 0.463639 | 0.500983 | day11 | [Atenolol] |
| I:P:C | 7.224887 | 29.2945 | 0.246629 | 0.806821 | 1 | 0.009812 | 0.009812 | 3.05E-05 | 0.99563 | day11 | [Atenolol] |
| I:E:C | -1.6621 | 30.19608 | -0.05504 | 0.956457 | 1 | 78.90727 | 78.90727 | 0.245196 | 0.62397 | day11 | [Atenolol] |
| P:E:C | 32.86568 | 30.19608 | 1.088409 | 0.284804 | 1 | 449.5849 | 449.5849 | 1.397038 | 0.246208 | day11 | [Atenolol] |
| I:P:E:C | -15.5564 | 42.07103 | -0.36976 | 0.714072 | 1 | 44.0001 | 44.0001 | 0.136726 | 0.714072 | day11 | [Atenolol] |

Supplementary Table 2: Statistic analysis of the microbial community databased on pharmaceutical combination. A linear model was also used to analyze how microbial biomass, microbial diversity, and the relative abundance of three taxa depended on the five compounds and their interactions. There was again one model for each of these five response variables. There were five binary explanatory variables, one for each of the five compounds. All possible two-, three-, four- and five-way interactions were included.

| **treatment_combination** | **estimate** | **std.error** | **t_statistic** | **t_p.value** | **df** | **sumsq** | **meansq** | **f_statistic** | **f_p.value** | **column** |
| --- | --- | --- | --- | --- | --- | --- | --- | --- | --- | --- |
| I | 10.66667 | 7.992831 | 1.334529 | 0.186759 | 1 | 175.2301 | 175.2301 | 1.828589 | 0.181053 | Biomass_day3 |
| P | 12.96667 | 7.992831 | 1.622287 | 0.109658 | 1 | 772.3676 | 772.3676 | 8.059935 | 0.006058 | Biomass_day3 |
| C | 13.7 | 7.992831 | 1.714036 | 0.091362 | 1 | 478.3801 | 478.3801 | 4.992069 | 0.028964 | Biomass_day3 |
| A | 15.76667 | 7.992831 | 1.972601 | 0.052864 | 1 | 1986.53 | 1986.53 | 20.73016 | 2.43E-05 | Biomass_day3 |
| E | 25.03333 | 7.992831 | 3.131973 | 0.002618 | 1 | 233.4384 | 233.4384 | 2.436014 | 0.123509 | Biomass_day3 |
| I:P | -6.66667 | 11.30357 | -0.58978 | 0.557412 | 1 | 64.1901 | 64.1901 | 0.669847 | 0.416142 | Biomass_day3 |
| I:C | -18.6333 | 11.30357 | -1.64845 | 0.104162 | 1 | 0.210938 | 0.210938 | 0.002201 | 0.962725 | Biomass_day3 |
| P:C | -1.53333 | 11.30357 | -0.13565 | 0.892523 | 1 | 7.877604 | 7.877604 | 0.082206 | 0.775257 | Biomass_day3 |
| I:A | -7.86667 | 11.30357 | -0.69595 | 0.488983 | 1 | 200.3926 | 200.3926 | 2.091169 | 0.153032 | Biomass_day3 |
| P:A | 2.6 | 11.30357 | 0.230016 | 0.818813 | 1 | 141.8634 | 141.8634 | 1.480396 | 0.228181 | Biomass_day3 |
| C:A | -8.96667 | 11.30357 | -0.79326 | 0.430558 | 1 | 50.89594 | 50.89594 | 0.531117 | 0.468796 | Biomass_day3 |
| I:E | -26.5667 | 11.30357 | -2.35029 | 0.021853 | 1 | 3.190104 | 3.190104 | 0.03329 | 0.855802 | Biomass_day3 |
| P:E | -18.8 | 11.30357 | -1.66319 | 0.101164 | 1 | 603.5051 | 603.5051 | 6.297794 | 0.01463 | Biomass_day3 |
| C:E | -14.3667 | 11.30357 | -1.27098 | 0.208334 | 1 | 0.017604 | 0.017604 | 0.000184 | 0.989228 | Biomass_day3 |
| A:E | -22.6333 | 11.30357 | -2.00232 | 0.049492 | 1 | 39.39844 | 39.39844 | 0.411137 | 0.523683 | Biomass_day3 |
| I:P:C | 4.466667 | 15.98566 | 0.279417 | 0.780826 | 1 | 4.995938 | 4.995938 | 0.052134 | 0.820118 | Biomass_day3 |
| I:P:A | -9.83333 | 15.98566 | -0.61513 | 0.540647 | 1 | 47.1801 | 47.1801 | 0.492341 | 0.485429 | Biomass_day3 |
| I:C:A | 9.3 | 15.98566 | 0.581771 | 0.562764 | 1 | 265.0026 | 265.0026 | 2.765398 | 0.101213 | Biomass_day3 |
| P:C:A | -2.93333 | 15.98566 | -0.1835 | 0.854988 | 1 | 43.0676 | 43.0676 | 0.449426 | 0.50502 | Biomass_day3 |
| I:P:E | 15.1 | 15.98566 | 0.944596 | 0.348417 | 1 | 4.725937 | 4.725937 | 0.049317 | 0.824963 | Biomass_day3 |
| I:C:E | 26.3 | 15.98566 | 1.645224 | 0.104827 | 1 | 243.5251 | 243.5251 | 2.541272 | 0.115832 | Biomass_day3 |
| P:C:E | 2.7 | 15.98566 | 0.168901 | 0.866407 | 1 | 64.5176 | 64.5176 | 0.673264 | 0.414962 | Biomass_day3 |
| I:A:E | 24.13333 | 15.98566 | 1.509686 | 0.136045 | 1 | 456.3176 | 456.3176 | 4.761839 | 0.032775 | Biomass_day3 |
| P:A:E | 9.3 | 15.98566 | 0.581771 | 0.562764 | 1 | 71.5876 | 71.5876 | 0.747042 | 0.390642 | Biomass_day3 |
| C:A:E | 9.633333 | 15.98566 | 0.602623 | 0.548889 | 1 | 73.6751 | 73.6751 | 0.768826 | 0.38386 | Biomass_day3 |
| I:P:C:A | 20.76667 | 22.60714 | 0.918589 | 0.36176 | 1 | 30.71344 | 30.71344 | 0.320506 | 0.573284 | Biomass_day3 |
| I:P:C:E | -14.3333 | 22.60714 | -0.63402 | 0.528329 | 1 | 254.4759 | 254.4759 | 2.655548 | 0.108102 | Biomass_day3 |
| I:P:A:E | -0.6 | 22.60714 | -0.02654 | 0.978909 | 1 | 56.8876 | 56.8876 | 0.593643 | 0.443849 | Biomass_day3 |
| I:C:A:E | -1.06667 | 22.60714 | -0.04718 | 0.962515 | 1 | 61.2801 | 61.2801 | 0.63948 | 0.426857 | Biomass_day3 |
| P:C:A:E | 7.533333 | 22.60714 | 0.333228 | 0.740051 | 1 | 6.562604 | 6.562604 | 0.068483 | 0.794399 | Biomass_day3 |
| I:P:C:A:E | -23.4333 | 31.97132 | -0.73295 | 0.466266 | 1 | 51.4801 | 51.4801 | 0.537213 | 0.466266 | Biomass_day3 |
| I | -6 | 5.614416 | -1.06868 | 0.289229 | 1 | 68.34375 | 68.34375 | 1.445434 | 0.233691 | Biomass_day11 |
| P | -12.1667 | 5.614416 | -2.16704 | 0.03396 | 1 | 137.2817 | 137.2817 | 2.903435 | 0.093242 | Biomass_day11 |
| C | 6.6 | 5.614416 | 1.175545 | 0.244131 | 1 | 85.88167 | 85.88167 | 1.816352 | 0.1825 | Biomass_day11 |
| A | -5.96667 | 5.614416 | -1.06274 | 0.291893 | 1 | 74.55375 | 74.55375 | 1.576773 | 0.213791 | Biomass_day11 |
| E | -12.6667 | 5.614416 | -2.2561 | 0.027489 | 1 | 239.4017 | 239.4017 | 5.063219 | 0.027884 | Biomass_day11 |
| I:P | 24.36667 | 7.939983 | 3.068856 | 0.00315 | 1 | 0.735 | 0.735 | 0.015545 | 0.901168 | Biomass_day11 |
| I:C | 1.166667 | 7.939983 | 0.146936 | 0.883645 | 1 | 11.20667 | 11.20667 | 0.237015 | 0.628034 | Biomass_day11 |
| P:C | -1.1 | 7.939983 | -0.13854 | 0.890249 | 1 | 91.65042 | 91.65042 | 1.938358 | 0.168666 | Biomass_day11 |
| I:A | 5.266667 | 7.939983 | 0.66331 | 0.509515 | 1 | 80.30042 | 80.30042 | 1.698312 | 0.197178 | Biomass_day11 |
| P:A | 14.5 | 7.939983 | 1.8262 | 0.072485 | 1 | 0.041667 | 0.041667 | 0.000881 | 0.97641 | Biomass_day11 |
| C:A | -5.2 | 7.939983 | -0.65491 | 0.514871 | 1 | 13.80167 | 13.80167 | 0.291898 | 0.590882 | Biomass_day11 |
| I:E | 15.4 | 7.939983 | 1.939551 | 0.056844 | 1 | 18.02667 | 18.02667 | 0.381255 | 0.539123 | Biomass_day11 |
| P:E | 16.26667 | 7.939983 | 2.048703 | 0.044597 | 1 | 88.55042 | 88.55042 | 1.872795 | 0.17594 | Biomass_day11 |
| C:E | 0.666667 | 7.939983 | 0.083963 | 0.933348 | 1 | 9.500417 | 9.500417 | 0.200929 | 0.655486 | Biomass_day11 |
| A:E | 12.23333 | 7.939983 | 1.540725 | 0.128314 | 1 | 36.50667 | 36.50667 | 0.772097 | 0.382856 | Biomass_day11 |
| I:P:C | -17.2 | 11.22883 | -1.53177 | 0.130507 | 1 | 39.78375 | 39.78375 | 0.841405 | 0.362439 | Biomass_day11 |
| I:P:A | -28.4 | 11.22883 | -2.5292 | 0.013907 | 1 | 319.74 | 319.74 | 6.762333 | 0.011549 | Biomass_day11 |
| I:C:A | 9.666667 | 11.22883 | 0.860879 | 0.392518 | 1 | 78.48167 | 78.48167 | 1.659846 | 0.202262 | Biomass_day11 |
| P:C:A | -0.13333 | 11.22883 | -0.01187 | 0.990563 | 1 | 10.80042 | 10.80042 | 0.228423 | 0.634324 | Biomass_day11 |
| I:P:E | -31.5 | 11.22883 | -2.80528 | 0.006651 | 1 | 44.55375 | 44.55375 | 0.942288 | 0.335344 | Biomass_day11 |
| I:C:E | -8.96667 | 11.22883 | -0.79854 | 0.427511 | 1 | 1.08375 | 1.08375 | 0.022921 | 0.88014 | Biomass_day11 |
| P:C:E | -4.96667 | 11.22883 | -0.44231 | 0.659753 | 1 | 153.015 | 153.015 | 3.236187 | 0.076743 | Biomass_day11 |
| I:A:E | -12.3 | 11.22883 | -1.09539 | 0.277449 | 1 | 0.806667 | 0.806667 | 0.017061 | 0.896489 | Biomass_day11 |
| P:A:E | -19.0667 | 11.22883 | -1.69801 | 0.094364 | 1 | 7.59375 | 7.59375 | 0.160604 | 0.689936 | Biomass_day11 |
| C:A:E | -4.36667 | 11.22883 | -0.38888 | 0.698656 | 1 | 21.47042 | 21.47042 | 0.454088 | 0.502827 | Biomass_day11 |
| I:P:C:A | 3.233333 | 15.87997 | 0.203611 | 0.839304 | 1 | 4.950417 | 4.950417 | 0.104699 | 0.747317 | Biomass_day11 |
| I:P:C:E | 27.73333 | 15.87997 | 1.746435 | 0.085533 | 1 | 163.2817 | 163.2817 | 3.453321 | 0.067725 | Biomass_day11 |
| I:P:A:E | 31.23333 | 15.87997 | 1.966839 | 0.053541 | 1 | 222.6504 | 222.6504 | 4.708939 | 0.033725 | Biomass_day11 |
| I:C:A:E | -1.23333 | 15.87997 | -0.07767 | 0.938336 | 1 | 24.60375 | 24.60375 | 0.520356 | 0.473317 | Biomass_day11 |
| P:C:A:E | 9.266667 | 15.87997 | 0.583544 | 0.561578 | 1 | 2.16 | 2.16 | 0.045683 | 0.831433 | Biomass_day11 |
| I:P:C:A:E | -13.7333 | 22.45766 | -0.61152 | 0.543021 | 1 | 17.68167 | 17.68167 | 0.373958 | 0.543021 | Biomass_day11 |
| I | -0.16323 | 0.37086 | -0.44015 | 0.661552 | 1 | 0.24773 | 0.24773 | 2.401573 | 0.126949 | Diversity_day3 |
| P | 0.00085 | 0.37086 | 0.002292 | 0.998179 | 1 | 1.933064 | 1.933064 | 18.73975 | 6.39E-05 | Diversity_day3 |
| C | 0.15454 | 0.37086 | 0.416707 | 0.678515 | 1 | 0.015132 | 0.015132 | 0.146698 | 0.703187 | Diversity_day3 |
| A | 0.483049 | 0.37086 | 1.302509 | 0.19817 | 1 | 0.065133 | 0.065133 | 0.631419 | 0.430251 | Diversity_day3 |
| E | -0.08547 | 0.37086 | -0.23048 | 0.818578 | 1 | 0.012728 | 0.012728 | 0.123393 | 0.726725 | Diversity_day3 |
| I:P | 0.243211 | 0.454209 | 0.53546 | 0.594491 | 1 | 1.250027 | 1.250027 | 12.11816 | 0.000986 | Diversity_day3 |
| I:C | -0.06291 | 0.454209 | -0.13851 | 0.890342 | 1 | 0.083936 | 0.083936 | 0.813702 | 0.370962 | Diversity_day3 |
| P:C | -0.39309 | 0.454209 | -0.86543 | 0.390563 | 1 | 0.134566 | 0.134566 | 1.304529 | 0.258336 | Diversity_day3 |
| I:A | 0.259818 | 0.472756 | 0.549582 | 0.584831 | 1 | 0.002822 | 0.002822 | 0.027358 | 0.869233 | Diversity_day3 |
| P:A | -1.32099 | 0.454209 | -2.90834 | 0.005232 | 1 | 1.757985 | 1.757985 | 17.04248 | 0.000125 | Diversity_day3 |
| C:A | -0.06926 | 0.472756 | -0.14649 | 0.884067 | 1 | 0.054725 | 0.054725 | 0.530527 | 0.469475 | Diversity_day3 |
| I:E | -0.02575 | 0.454209 | -0.05669 | 0.955 | 1 | 0.138149 | 0.138149 | 1.33926 | 0.252166 | Diversity_day3 |
| P:E | -0.1748 | 0.524476 | -0.33328 | 0.740192 | 1 | 1.027572 | 1.027572 | 9.961621 | 0.002593 | Diversity_day3 |
| C:E | 0.391831 | 0.472756 | 0.828823 | 0.410787 | 1 | 0.109801 | 0.109801 | 1.064445 | 0.306719 | Diversity_day3 |
| A:E | -0.0777 | 0.472756 | -0.16436 | 0.87005 | 1 | 0.02403 | 0.02403 | 0.232955 | 0.631257 | Diversity_day3 |
| I:P:C | 0.092422 | 0.586382 | 0.157614 | 0.875338 | 1 | 0.000672 | 0.000672 | 0.006514 | 0.935966 | Diversity_day3 |
| I:P:A | 0.00953 | 0.600863 | 0.01586 | 0.987404 | 1 | 0.092387 | 0.092387 | 0.895634 | 0.348095 | Diversity_day3 |
| I:C:A | -0.52206 | 0.615003 | -0.84887 | 0.399634 | 1 | 0.003605 | 0.003605 | 0.034946 | 0.852397 | Diversity_day3 |
| P:C:A | 0.671747 | 0.600863 | 1.11797 | 0.268439 | 1 | 0.008411 | 0.008411 | 0.08154 | 0.776294 | Diversity_day3 |
| I:P:E | 0.083153 | 0.642349 | 0.129452 | 0.897472 | 1 | 0.062781 | 0.062781 | 0.608619 | 0.438653 | Diversity_day3 |
| I:C:E | -0.42757 | 0.600863 | -0.7116 | 0.479719 | 1 | 0.055058 | 0.055058 | 0.533748 | 0.468136 | Diversity_day3 |
| P:C:E | 0.217041 | 0.655595 | 0.33106 | 0.741857 | 1 | 0.168711 | 0.168711 | 1.635539 | 0.206308 | Diversity_day3 |
| I:A:E | -0.82547 | 0.615003 | -1.34223 | 0.185037 | 1 | 0.005674 | 0.005674 | 0.055009 | 0.815437 | Diversity_day3 |
| P:A:E | 1.060794 | 0.655595 | 1.618064 | 0.11137 | 1 | 0.743139 | 0.743139 | 7.204226 | 0.009591 | Diversity_day3 |
| C:A:E | -0.42609 | 0.642349 | -0.66334 | 0.509886 | 1 | 0.102771 | 0.102771 | 0.9963 | 0.322579 | Diversity_day3 |
| I:P:C:A | 0.022653 | 0.808272 | 0.028027 | 0.977742 | 1 | 0.062677 | 0.062677 | 0.607616 | 0.439028 | Diversity_day3 |
| I:P:C:E | 0.139017 | 0.839571 | 0.165581 | 0.869093 | 1 | 0.002114 | 0.002114 | 0.020493 | 0.886692 | Diversity_day3 |
| I:P:A:E | 0.629359 | 0.849748 | 0.740641 | 0.462061 | 1 | 0.042346 | 0.042346 | 0.410519 | 0.524367 | Diversity_day3 |
| I:C:A:E | 1.38516 | 0.839571 | 1.649842 | 0.104676 | 1 | 0.366223 | 0.366223 | 3.550283 | 0.064825 | Diversity_day3 |
| P:C:A:E | -0.8298 | 0.869745 | -0.95407 | 0.344221 | 1 | 0.454279 | 0.454279 | 4.403933 | 0.040462 | Diversity_day3 |
| I:P:C:A:E | -0.63889 | 1.150564 | -0.55528 | 0.580954 | 1 | 0.031806 | 0.031806 | 0.308337 | 0.580954 | Diversity_day3 |
| I | 1.16692 | 0.243411 | 4.79404 | 1.04E-05 | 1 | 0.617978 | 0.617978 | 6.953484 | 0.010523 | Diversity_day11 |
| P | 0.602657 | 0.243411 | 2.475884 | 0.01599 | 1 | 38.70512 | 38.70512 | 435.5098 | 5.39E-30 | Diversity_day11 |
| C | 1.121457 | 0.272141 | 4.120861 | 0.000112 | 1 | 0.324184 | 0.324184 | 3.647713 | 0.060701 | Diversity_day11 |
| A | 2.017197 | 0.243411 | 8.287217 | 1.12E-11 | 1 | 1.055887 | 1.055887 | 11.88084 | 0.001015 | Diversity_day11 |
| E | 0.825518 | 0.243411 | 3.391464 | 0.001205 | 1 | 0.278587 | 0.278587 | 3.134663 | 0.081482 | Diversity_day11 |
| I:P | -1.55602 | 0.344235 | -4.52024 | 2.78E-05 | 1 | 0.050933 | 0.050933 | 0.573095 | 0.451854 | Diversity_day11 |
| I:C | -1.13486 | 0.365116 | -3.10822 | 0.002824 | 1 | 0.312276 | 0.312276 | 3.51373 | 0.0655 | Diversity_day11 |
| P:C | -1.7581 | 0.365116 | -4.81519 | 9.59E-06 | 1 | 1.107258 | 1.107258 | 12.45886 | 0.000783 | Diversity_day11 |
| I:A | -1.21175 | 0.344235 | -3.52014 | 0.000807 | 1 | 0.295554 | 0.295554 | 3.325577 | 0.072954 | Diversity_day11 |
| P:A | -2.3578 | 0.344235 | -6.84941 | 3.65E-09 | 1 | 7.030357 | 7.030357 | 79.10554 | 9.88E-13 | Diversity_day11 |
| C:A | -1.39863 | 0.365116 | -3.83064 | 0.000297 | 1 | 1.001322 | 1.001322 | 11.26687 | 0.001341 | Diversity_day11 |
| I:E | -1.37668 | 0.344235 | -3.99924 | 0.000169 | 1 | 0.493583 | 0.493583 | 5.553798 | 0.021563 | Diversity_day11 |
| P:E | -1.16059 | 0.344235 | -3.37152 | 0.001282 | 1 | 0.009479 | 0.009479 | 0.10666 | 0.745062 | Diversity_day11 |
| C:E | -0.36274 | 0.365116 | -0.9935 | 0.324266 | 1 | 0.139224 | 0.139224 | 1.566551 | 0.215336 | Diversity_day11 |
| A:E | -1.01928 | 0.344235 | -2.96101 | 0.004319 | 1 | 0.237031 | 0.237031 | 2.667071 | 0.107433 | Diversity_day11 |
| I:P:C | 1.569963 | 0.501804 | 3.128637 | 0.002659 | 1 | 0.343259 | 0.343259 | 3.862347 | 0.053794 | Diversity_day11 |
| I:P:A | 1.35906 | 0.486821 | 2.791702 | 0.006932 | 1 | 0.010415 | 0.010415 | 0.117185 | 0.733248 | Diversity_day11 |
| I:C:A | 0.911955 | 0.501804 | 1.817354 | 0.073919 | 1 | 0.084048 | 0.084048 | 0.945712 | 0.334532 | Diversity_day11 |
| P:C:A | 1.644158 | 0.501804 | 3.276495 | 0.001712 | 1 | 0.786608 | 0.786608 | 8.850904 | 0.00415 | Diversity_day11 |
| I:P:E | 1.670244 | 0.486821 | 3.430918 | 0.001067 | 1 | 0.959029 | 0.959029 | 10.79098 | 0.001668 | Diversity_day11 |
| I:C:E | 0.410724 | 0.501804 | 0.818496 | 0.416159 | 1 | 0.033158 | 0.033158 | 0.373096 | 0.543519 | Diversity_day11 |
| P:C:E | 0.780597 | 0.501804 | 1.555581 | 0.124817 | 1 | 0.109658 | 0.109658 | 1.233867 | 0.270878 | Diversity_day11 |
| I:A:E | 0.95584 | 0.486821 | 1.963431 | 0.054013 | 1 | 0.376715 | 0.376715 | 4.238799 | 0.04365 | Diversity_day11 |
| P:A:E | 1.060905 | 0.486821 | 2.17925 | 0.03306 | 1 | 0.195684 | 0.195684 | 2.201832 | 0.142833 | Diversity_day11 |
| C:A:E | 0.328211 | 0.501804 | 0.654062 | 0.515452 | 1 | 0.074136 | 0.074136 | 0.834175 | 0.36455 | Diversity_day11 |
| I:P:C:A | -1.62077 | 0.699144 | -2.31822 | 0.023697 | 1 | 0.694726 | 0.694726 | 7.817054 | 0.006853 | Diversity_day11 |
| I:P:C:E | -0.79718 | 0.699144 | -1.14022 | 0.258511 | 1 | 0.113994 | 0.113994 | 1.282663 | 0.261697 | Diversity_day11 |
| I:P:A:E | -1.19665 | 0.688469 | -1.73813 | 0.087074 | 1 | 0.331125 | 0.331125 | 3.725811 | 0.058082 | Diversity_day11 |
| I:C:A:E | 0.028558 | 0.699144 | 0.040848 | 0.967546 | 1 | 0.030763 | 0.030763 | 0.346143 | 0.558408 | Diversity_day11 |
| P:C:A:E | -0.47345 | 0.699144 | -0.67719 | 0.500765 | 1 | 0.017346 | 0.017346 | 0.195172 | 0.660159 | Diversity_day11 |
| I:P:C:A:E | 0.505692 | 0.98122 | 0.51537 | 0.608098 | 1 | 0.023605 | 0.023605 | 0.265607 | 0.608098 | Diversity_day11 |
| I | 0.006401 | 0.018951 | 0.337747 | 0.73684 | 1 | 0.009849 | 0.009849 | 36.56415 | 1.35E-07 | Achromobacter_day3 |
| P | 0.015181 | 0.018951 | 0.801092 | 0.426525 | 1 | 0.007502 | 0.007502 | 27.85074 | 2.29E-06 | Achromobacter_day3 |
| C | -0.006 | 0.018951 | -0.31641 | 0.752885 | 1 | 0.000137 | 0.000137 | 0.510275 | 0.478041 | Achromobacter_day3 |
| A | -0.00399 | 0.018951 | -0.21053 | 0.834028 | 1 | 0.004202 | 0.004202 | 15.59869 | 0.000225 | Achromobacter_day3 |
| E | -0.00554 | 0.018951 | -0.2923 | 0.771159 | 1 | 0.003336 | 0.003336 | 12.38391 | 0.000878 | Achromobacter_day3 |
| I:P | 0.029297 | 0.02321 | 1.262265 | 0.21218 | 1 | 0.002107 | 0.002107 | 7.821651 | 0.007099 | Achromobacter_day3 |
| I:C | 0.011757 | 0.02321 | 0.506533 | 0.614507 | 1 | 9.65E-05 | 9.65E-05 | 0.358367 | 0.551872 | Achromobacter_day3 |
| P:C | -0.00306 | 0.02321 | -0.13202 | 0.89545 | 1 | 0.000144 | 0.000144 | 0.535203 | 0.467533 | Achromobacter_day3 |
| I:A | 0.007901 | 0.024158 | 0.327072 | 0.744855 | 1 | 0.000398 | 0.000398 | 1.476127 | 0.229572 | Achromobacter_day3 |
| P:A | -0.026 | 0.02321 | -1.12 | 0.267582 | 1 | 0.001933 | 0.001933 | 7.176897 | 0.009721 | Achromobacter_day3 |
| C:A | 0.006259 | 0.024158 | 0.259104 | 0.796522 | 1 | 9.34E-05 | 9.34E-05 | 0.346729 | 0.558381 | Achromobacter_day3 |
| I:E | 0.01053 | 0.02321 | 0.453676 | 0.651847 | 1 | 0.000341 | 0.000341 | 1.26673 | 0.265268 | Achromobacter_day3 |
| P:E | -0.00065 | 0.026801 | -0.02412 | 0.980844 | 1 | 0.002153 | 0.002153 | 7.994302 | 0.006532 | Achromobacter_day3 |
| C:E | 0.015565 | 0.024158 | 0.644288 | 0.522067 | 1 | 0.000314 | 0.000314 | 1.16529 | 0.285082 | Achromobacter_day3 |
| A:E | 0.006149 | 0.024158 | 0.254553 | 0.800018 | 1 | 0.000895 | 0.000895 | 3.321973 | 0.073797 | Achromobacter_day3 |
| I:P:C | -0.01294 | 0.029964 | -0.43181 | 0.66757 | 1 | 6.53E-05 | 6.53E-05 | 0.242414 | 0.624428 | Achromobacter_day3 |
| I:P:A | -0.0347 | 0.030704 | -1.12999 | 0.263384 | 1 | 8.41E-06 | 8.41E-06 | 0.031216 | 0.860408 | Achromobacter_day3 |
| I:C:A | -0.02087 | 0.031427 | -0.66421 | 0.509329 | 1 | 9.89E-05 | 9.89E-05 | 0.367248 | 0.547001 | Achromobacter_day3 |
| P:C:A | 0.00839 | 0.030704 | 0.273268 | 0.785671 | 1 | 0.000193 | 0.000193 | 0.715214 | 0.401383 | Achromobacter_day3 |
| I:P:E | -0.01596 | 0.032824 | -0.48629 | 0.62869 | 1 | 0.000389 | 0.000389 | 1.444189 | 0.234612 | Achromobacter_day3 |
| I:C:E | -0.01485 | 0.030704 | -0.48352 | 0.630646 | 1 | 2.16E-05 | 2.16E-05 | 0.080317 | 0.777932 | Achromobacter_day3 |
| P:C:E | 0.006052 | 0.033501 | 0.180656 | 0.857302 | 1 | 7.01E-05 | 7.01E-05 | 0.260428 | 0.611868 | Achromobacter_day3 |
| I:A:E | -0.02974 | 0.031427 | -0.94649 | 0.348038 | 1 | 0.000255 | 0.000255 | 0.945816 | 0.335045 | Achromobacter_day3 |
| P:A:E | 0.023162 | 0.033501 | 0.691377 | 0.492237 | 1 | 0.001536 | 0.001536 | 5.702971 | 0.020397 | Achromobacter_day3 |
| C:A:E | -0.01421 | 0.032824 | -0.43306 | 0.666666 | 1 | 0.000581 | 0.000581 | 2.156545 | 0.14766 | Achromobacter_day3 |
| I:P:C:A | 0.020577 | 0.041303 | 0.498205 | 0.620326 | 1 | 0.000123 | 0.000123 | 0.457903 | 0.501442 | Achromobacter_day3 |
| I:P:C:E | 0.031482 | 0.042902 | 0.733802 | 0.466186 | 1 | 1.04E-05 | 1.04E-05 | 0.038782 | 0.844607 | Achromobacter_day3 |
| I:P:A:E | 0.076553 | 0.043422 | 1.762997 | 0.083456 | 1 | 0.000492 | 0.000492 | 1.828395 | 0.181851 | Achromobacter_day3 |
| I:C:A:E | 0.045415 | 0.042902 | 1.058585 | 0.294416 | 1 | 1.91E-05 | 1.91E-05 | 0.070808 | 0.791159 | Achromobacter_day3 |
| P:C:A:E | -0.01854 | 0.044444 | -0.41725 | 0.678122 | 1 | 0.00118 | 0.00118 | 4.379615 | 0.040998 | Achromobacter_day3 |
| I:P:C:A:E | -0.0741 | 0.058794 | -1.26041 | 0.212842 | 1 | 0.000428 | 0.000428 | 1.588638 | 0.212842 | Achromobacter_day3 |
| I | -0.34653 | 0.070219 | -4.93507 | 6.18E-06 | 1 | 0.012657 | 0.012657 | 1.711369 | 0.195561 | Achromobacter_day11 |
| P | 0.009274 | 0.070219 | 0.132075 | 0.895346 | 1 | 3.960783 | 3.960783 | 535.5324 | 1.68E-32 | Achromobacter_day11 |
| C | -0.36657 | 0.078507 | -4.66929 | 1.63E-05 | 1 | 6.41E-06 | 6.41E-06 | 0.000866 | 0.976615 | Achromobacter_day11 |
| A | -0.36396 | 0.070219 | -5.1832 | 2.45E-06 | 1 | 0.000297 | 0.000297 | 0.040134 | 0.841864 | Achromobacter_day11 |
| E | -0.37794 | 0.070219 | -5.38232 | 1.15E-06 | 1 | 0.023915 | 0.023915 | 3.233521 | 0.076937 | Achromobacter_day11 |
| I:P | 0.427019 | 0.099304 | 4.300115 | 6.05E-05 | 1 | 0.009311 | 0.009311 | 1.258979 | 0.266103 | Achromobacter_day11 |
| I:C | 0.329061 | 0.105328 | 3.124163 | 0.002695 | 1 | 0.005511 | 0.005511 | 0.745191 | 0.391278 | Achromobacter_day11 |
| P:C | 0.306347 | 0.105328 | 2.908511 | 0.005011 | 1 | 0.026796 | 0.026796 | 3.623041 | 0.061555 | Achromobacter_day11 |
| I:A | 0.373988 | 0.099304 | 3.766086 | 0.000367 | 1 | 0.032184 | 0.032184 | 4.351516 | 0.041031 | Achromobacter_day11 |
| P:A | 0.327078 | 0.099304 | 3.293707 | 0.001625 | 1 | 0.022158 | 0.022158 | 2.996008 | 0.088365 | Achromobacter_day11 |
| C:A | 0.344328 | 0.105328 | 3.269103 | 0.00175 | 1 | 0.038458 | 0.038458 | 5.199803 | 0.025984 | Achromobacter_day11 |
| I:E | 0.333875 | 0.099304 | 3.362153 | 0.001319 | 1 | 0.000782 | 0.000782 | 0.105669 | 0.746207 | Achromobacter_day11 |
| P:E | 0.471959 | 0.099304 | 4.752661 | 1.21E-05 | 1 | 0.137901 | 0.137901 | 18.64542 | 5.68E-05 | Achromobacter_day11 |
| C:E | 0.410159 | 0.105328 | 3.89412 | 0.000241 | 1 | 0.074628 | 0.074628 | 10.09033 | 0.002309 | Achromobacter_day11 |
| A:E | 0.376605 | 0.099304 | 3.792444 | 0.000337 | 1 | 0.063155 | 0.063155 | 8.539051 | 0.004821 | Achromobacter_day11 |
| I:P:C | -0.30258 | 0.144759 | -2.0902 | 0.040641 | 1 | 0.032091 | 0.032091 | 4.338995 | 0.041313 | Achromobacter_day11 |
| I:P:A | -0.42756 | 0.140437 | -3.0445 | 0.003399 | 1 | 0.028108 | 0.028108 | 3.800436 | 0.055694 | Achromobacter_day11 |
| I:C:A | -0.32881 | 0.144759 | -2.2714 | 0.026548 | 1 | 0.010964 | 0.010964 | 1.482367 | 0.227945 | Achromobacter_day11 |
| P:C:A | -0.18255 | 0.144759 | -1.26109 | 0.211927 | 1 | 0.003778 | 0.003778 | 0.510763 | 0.477449 | Achromobacter_day11 |
| I:P:E | -0.50602 | 0.140437 | -3.60316 | 0.000621 | 1 | 0.03126 | 0.03126 | 4.22659 | 0.043945 | Achromobacter_day11 |
| I:C:E | -0.36534 | 0.144759 | -2.5238 | 0.014143 | 1 | 0.003842 | 0.003842 | 0.519498 | 0.473722 | Achromobacter_day11 |
| P:C:E | -0.28417 | 0.144759 | -1.96305 | 0.054059 | 1 | 0.009693 | 0.009693 | 1.310547 | 0.256626 | Achromobacter_day11 |
| I:A:E | -0.39184 | 0.140437 | -2.79016 | 0.006962 | 1 | 0.005004 | 0.005004 | 0.676631 | 0.413852 | Achromobacter_day11 |
| P:A:E | -0.36071 | 0.140437 | -2.56846 | 0.012597 | 1 | 0.000656 | 0.000656 | 0.088715 | 0.766798 | Achromobacter_day11 |
| C:A:E | -0.39277 | 0.144759 | -2.71323 | 0.008582 | 1 | 0.016235 | 0.016235 | 2.195147 | 0.143431 | Achromobacter_day11 |
| I:P:C:A | 0.184455 | 0.201688 | 0.914556 | 0.363912 | 1 | 0.000277 | 0.000277 | 0.037453 | 0.847169 | Achromobacter_day11 |
| I:P:C:E | 0.326565 | 0.201688 | 1.619162 | 0.110408 | 1 | 0.005009 | 0.005009 | 0.677284 | 0.413628 | Achromobacter_day11 |
| I:P:A:E | 0.613777 | 0.198608 | 3.090391 | 0.002975 | 1 | 0.063518 | 0.063518 | 8.588173 | 0.004708 | Achromobacter_day11 |
| I:C:A:E | 0.500031 | 0.201688 | 2.479237 | 0.015854 | 1 | 0.030659 | 0.030659 | 4.145398 | 0.045958 | Achromobacter_day11 |
| P:C:A:E | 0.279905 | 0.201688 | 1.387818 | 0.170082 | 1 | 0.001759 | 0.001759 | 0.237821 | 0.627477 | Achromobacter_day11 |
| I:P:C:A:E | -0.4154 | 0.28306 | -1.46752 | 0.147209 | 1 | 0.015928 | 0.015928 | 2.15362 | 0.147209 | Achromobacter_day11 |
| I | -0.0649 | 0.115057 | -0.56406 | 0.575006 | 1 | 0.471907 | 0.471907 | 47.53019 | 5.62E-09 | Acinetobacter_day3 |
| P | 0.200508 | 0.115057 | 1.742683 | 0.086978 | 1 | 0.292997 | 0.292997 | 29.51052 | 1.3E-06 | Acinetobacter_day3 |
| C | 0.06129 | 0.115057 | 0.532691 | 0.596394 | 1 | 0.126068 | 0.126068 | 12.69748 | 0.000766 | Acinetobacter_day3 |
| A | -0.01273 | 0.115057 | -0.11068 | 0.912271 | 1 | 0.048565 | 0.048565 | 4.891445 | 0.031164 | Acinetobacter_day3 |
| E | 0.148652 | 0.115057 | 1.29199 | 0.201763 | 1 | 0.052638 | 0.052638 | 5.301644 | 0.025114 | Acinetobacter_day3 |
| I:P | -0.1118 | 0.140915 | -0.79338 | 0.430969 | 1 | 0.041083 | 0.041083 | 4.137876 | 0.046768 | Acinetobacter_day3 |
| I:C | -0.12688 | 0.140915 | -0.90038 | 0.371845 | 1 | 0.000873 | 0.000873 | 0.08792 | 0.767954 | Acinetobacter_day3 |
| P:C | -0.05727 | 0.140915 | -0.4064 | 0.686025 | 1 | 0.001878 | 0.001878 | 0.189168 | 0.665312 | Acinetobacter_day3 |
| I:A | -0.00902 | 0.146669 | -0.06148 | 0.951196 | 1 | 0.08579 | 0.08579 | 8.640683 | 0.004799 | Acinetobacter_day3 |
| P:A | 0.164937 | 0.140915 | 1.170469 | 0.246859 | 1 | 0.08021 | 0.08021 | 8.078668 | 0.006273 | Acinetobacter_day3 |
| C:A | -0.11794 | 0.146669 | -0.80411 | 0.424795 | 1 | 0.001026 | 0.001026 | 0.103289 | 0.749136 | Acinetobacter_day3 |
| I:E | -0.14348 | 0.140915 | -1.01823 | 0.313026 | 1 | 0.008997 | 0.008997 | 0.906197 | 0.34529 | Acinetobacter_day3 |
| P:E | -0.0829 | 0.162715 | -0.50947 | 0.612464 | 1 | 0.120252 | 0.120252 | 12.11174 | 0.000988 | Acinetobacter_day3 |
| C:E | -0.16544 | 0.146669 | -1.12796 | 0.264235 | 1 | 0.012081 | 0.012081 | 1.216811 | 0.274791 | Acinetobacter_day3 |
| A:E | -0.08614 | 0.146669 | -0.58729 | 0.559411 | 1 | 0.011605 | 0.011605 | 1.168883 | 0.284349 | Acinetobacter_day3 |
| I:P:C | 0.028479 | 0.181921 | 0.156545 | 0.876177 | 1 | 0.001794 | 0.001794 | 0.180703 | 0.672429 | Acinetobacter_day3 |
| I:P:A | 0.078369 | 0.186414 | 0.420402 | 0.67583 | 1 | 0.003311 | 0.003311 | 0.333443 | 0.565995 | Acinetobacter_day3 |
| I:C:A | 0.250398 | 0.1908 | 1.312357 | 0.19485 | 1 | 8.24E-05 | 8.24E-05 | 0.008296 | 0.927759 | Acinetobacter_day3 |
| P:C:A | -0.04581 | 0.186414 | -0.24572 | 0.806815 | 1 | 0.001344 | 0.001344 | 0.135351 | 0.714359 | Acinetobacter_day3 |
| I:P:E | -0.01472 | 0.199284 | -0.07388 | 0.941375 | 1 | 0.00029 | 0.00029 | 0.029188 | 0.864974 | Acinetobacter_day3 |
| I:C:E | 0.17769 | 0.186414 | 0.953204 | 0.344657 | 1 | 6.63E-06 | 6.63E-06 | 0.000668 | 0.979475 | Acinetobacter_day3 |
| P:C:E | -0.0506 | 0.203394 | -0.24877 | 0.804466 | 1 | 0.021914 | 0.021914 | 2.207212 | 0.143077 | Acinetobacter_day3 |
| I:A:E | 0.253914 | 0.1908 | 1.330781 | 0.188752 | 1 | 0.002679 | 0.002679 | 0.269875 | 0.6055 | Acinetobacter_day3 |
| P:A:E | -0.15171 | 0.203394 | -0.7459 | 0.458906 | 1 | 0.008458 | 0.008458 | 0.851843 | 0.360063 | Acinetobacter_day3 |
| C:A:E | 0.139504 | 0.199284 | 0.700025 | 0.486862 | 1 | 0.001037 | 0.001037 | 0.104421 | 0.747812 | Acinetobacter_day3 |
| I:P:C:A | -0.09785 | 0.250761 | -0.39023 | 0.697876 | 1 | 4.85E-05 | 4.85E-05 | 0.004887 | 0.944522 | Acinetobacter_day3 |
| I:P:C:E | 0.063925 | 0.260471 | 0.245422 | 0.807043 | 1 | 0.002978 | 0.002978 | 0.299941 | 0.586135 | Acinetobacter_day3 |
| I:P:A:E | -0.1566 | 0.263629 | -0.59401 | 0.554943 | 1 | 0.004017 | 0.004017 | 0.404595 | 0.527364 | Acinetobacter_day3 |
| I:C:A:E | -0.45279 | 0.260471 | -1.73834 | 0.087747 | 1 | 0.049187 | 0.049187 | 4.954085 | 0.030147 | Acinetobacter_day3 |
| P:C:A:E | 0.205959 | 0.269832 | 0.763286 | 0.448554 | 1 | 0.02417 | 0.02417 | 2.434407 | 0.124435 | Acinetobacter_day3 |
| I:P:C:A:E | 0.121897 | 0.356955 | 0.341491 | 0.734036 | 1 | 0.001158 | 0.001158 | 0.116616 | 0.734036 | Acinetobacter_day3 |
| I | 0.053896 | 0.016562 | 3.254271 | 0.00183 | 1 | 0.000384 | 0.000384 | 0.932625 | 0.337874 | Acinetobacter_day11 |
| P | 0.087744 | 0.016562 | 5.297961 | 1.59E-06 | 1 | 0.015022 | 0.015022 | 36.51075 | 8.99E-08 | Acinetobacter_day11 |
| C | 0.017517 | 0.018517 | 0.946025 | 0.34775 | 1 | 0.021422 | 0.021422 | 52.06619 | 8.4E-10 | Acinetobacter_day11 |
| A | 0.012383 | 0.016562 | 0.747685 | 0.457432 | 1 | 0.003563 | 0.003563 | 8.660262 | 0.004548 | Acinetobacter_day11 |
| E | 0.022366 | 0.016562 | 1.35045 | 0.181703 | 1 | 0.003532 | 0.003532 | 8.58355 | 0.004719 | Acinetobacter_day11 |
| I:P | -0.07997 | 0.023422 | -3.4142 | 0.001124 | 1 | 0.008699 | 0.008699 | 21.14201 | 2.11E-05 | Acinetobacter_day11 |
| I:C | -0.0226 | 0.024843 | -0.90991 | 0.366336 | 1 | 0.000472 | 0.000472 | 1.148389 | 0.287975 | Acinetobacter_day11 |
| P:C | -0.06359 | 0.024843 | -2.55988 | 0.012882 | 1 | 0.0072 | 0.0072 | 17.50063 | 9.06E-05 | Acinetobacter_day11 |
| I:A | -0.04364 | 0.023422 | -1.86335 | 0.067075 | 1 | 0.000845 | 0.000845 | 2.053236 | 0.156827 | Acinetobacter_day11 |
| P:A | 0.03707 | 0.023422 | 1.582708 | 0.118496 | 1 | 0.001812 | 0.001812 | 4.403347 | 0.039884 | Acinetobacter_day11 |
| C:A | -0.02512 | 0.024843 | -1.01119 | 0.315793 | 1 | 0.000922 | 0.000922 | 2.241086 | 0.13938 | Acinetobacter_day11 |
| I:E | -0.01719 | 0.023422 | -0.73411 | 0.465605 | 1 | 0.000264 | 0.000264 | 0.640969 | 0.426369 | Acinetobacter_day11 |
| P:E | -0.02876 | 0.023422 | -1.22802 | 0.224006 | 1 | 0.004185 | 0.004185 | 10.17089 | 0.002224 | Acinetobacter_day11 |
| C:E | -0.00726 | 0.024843 | -0.29219 | 0.771098 | 1 | 0.000131 | 0.000131 | 0.318502 | 0.574513 | Acinetobacter_day11 |
| A:E | -0.02289 | 0.023422 | -0.9773 | 0.332159 | 1 | 0.001298 | 0.001298 | 3.154374 | 0.080553 | Acinetobacter_day11 |
| I:P:C | 0.029841 | 0.034143 | 0.873997 | 0.385439 | 1 | 0.00261 | 0.00261 | 6.344624 | 0.014325 | Acinetobacter_day11 |
| I:P:A | 3.25E-05 | 0.033124 | 0.000981 | 0.99922 | 1 | 0.000481 | 0.000481 | 1.168951 | 0.283738 | Acinetobacter_day11 |
| I:C:A | 0.007292 | 0.034143 | 0.213575 | 0.831568 | 1 | 2.6E-07 | 2.6E-07 | 0.000633 | 0.98001 | Acinetobacter_day11 |
| P:C:A | -0.02609 | 0.034143 | -0.76405 | 0.44769 | 1 | 1.96E-05 | 1.96E-05 | 0.047695 | 0.827829 | Acinetobacter_day11 |
| I:P:E | 0.00587 | 0.033124 | 0.177226 | 0.859899 | 1 | 0.002596 | 0.002596 | 6.308637 | 0.014591 | Acinetobacter_day11 |
| I:C:E | -0.0326 | 0.034143 | -0.95484 | 0.343306 | 1 | 0.000826 | 0.000826 | 2.007595 | 0.16144 | Acinetobacter_day11 |
| P:C:E | -0.00568 | 0.034143 | -0.16623 | 0.868512 | 1 | 0.002518 | 0.002518 | 6.119832 | 0.016074 | Acinetobacter_day11 |
| I:A:E | 0.02738 | 0.033124 | 0.826591 | 0.411589 | 1 | 0.002881 | 0.002881 | 7.002347 | 0.010268 | Acinetobacter_day11 |
| P:A:E | -0.07964 | 0.033124 | -2.40432 | 0.019155 | 1 | 0.000989 | 0.000989 | 2.403762 | 0.126053 | Acinetobacter_day11 |
| C:A:E | 0.004709 | 0.034143 | 0.137926 | 0.890739 | 1 | 0.001536 | 0.001536 | 3.733035 | 0.057846 | Acinetobacter_day11 |
| I:P:C:A | 0.025345 | 0.04757 | 0.532803 | 0.596045 | 1 | 0.000516 | 0.000516 | 1.253272 | 0.267178 | Acinetobacter_day11 |
| I:P:C:E | 0.060359 | 0.04757 | 1.268854 | 0.209161 | 1 | 2.08E-06 | 2.08E-06 | 0.005044 | 0.943604 | Acinetobacter_day11 |
| I:P:A:E | 0.072908 | 0.046844 | 1.556416 | 0.124619 | 1 | 5.01E-05 | 5.01E-05 | 0.121848 | 0.728203 | Acinetobacter_day11 |
| I:C:A:E | 0.021126 | 0.04757 | 0.444094 | 0.658497 | 1 | 0.000664 | 0.000664 | 1.614691 | 0.208506 | Acinetobacter_day11 |
| P:C:A:E | 0.095909 | 0.04757 | 2.016164 | 0.048051 | 1 | 0.000399 | 0.000399 | 0.970046 | 0.328436 | Acinetobacter_day11 |
| I:P:C:A:E | -0.12416 | 0.066763 | -1.85973 | 0.067594 | 1 | 0.001423 | 0.001423 | 3.458604 | 0.067594 | Acinetobacter_day11 |
| I | -0.02939 | 0.0292 | -1.00656 | 0.318553 | 1 | 0.138124 | 0.138124 | 215.9935 | 1.07E-20 | Comamonas_day3 |
| P | -0.00691 | 0.0292 | -0.23673 | 0.813746 | 1 | 0.018346 | 0.018346 | 28.68936 | 1.72E-06 | Comamonas_day3 |
| C | 0.075263 | 0.0292 | 2.577515 | 0.012662 | 1 | 0.000248 | 0.000248 | 0.388048 | 0.535902 | Comamonas_day3 |
| A | 0.087961 | 0.0292 | 3.012378 | 0.003912 | 1 | 0.002457 | 0.002457 | 3.841682 | 0.055068 | Comamonas_day3 |
| E | 0.02823 | 0.0292 | 0.966764 | 0.337894 | 1 | 0.001187 | 0.001187 | 1.85628 | 0.178608 | Comamonas_day3 |
| I:P | -0.01212 | 0.035763 | -0.33893 | 0.735951 | 1 | 0.007132 | 0.007132 | 11.15225 | 0.001512 | Comamonas_day3 |
| I:C | -0.09512 | 0.035763 | -2.65985 | 0.010222 | 1 | 0.001074 | 0.001074 | 1.679291 | 0.200431 | Comamonas_day3 |
| P:C | -0.06357 | 0.035763 | -1.77753 | 0.08101 | 1 | 4.48E-05 | 4.48E-05 | 0.070126 | 0.792144 | Comamonas_day3 |
| I:A | -0.07062 | 0.037223 | -1.8972 | 0.063057 | 1 | 3.34E-05 | 3.34E-05 | 0.052221 | 0.82009 | Comamonas_day3 |
| P:A | -0.05962 | 0.035763 | -1.66703 | 0.101193 | 1 | 0.001692 | 0.001692 | 2.645222 | 0.109578 | Comamonas_day3 |
| C:A | -0.06857 | 0.037223 | -1.84224 | 0.070833 | 1 | 0.001097 | 0.001097 | 1.715942 | 0.195661 | Comamonas_day3 |
| I:E | -0.04571 | 0.035763 | -1.2782 | 0.206548 | 1 | 0.002459 | 0.002459 | 3.845417 | 0.054953 | Comamonas_day3 |
| P:E | 0.050705 | 0.041295 | 1.227863 | 0.224727 | 1 | 0.003172 | 0.003172 | 4.959692 | 0.030057 | Comamonas_day3 |
| C:E | -0.05559 | 0.037223 | -1.49337 | 0.141056 | 1 | 0.000118 | 0.000118 | 0.184523 | 0.669193 | Comamonas_day3 |
| A:E | -0.03572 | 0.037223 | -0.95964 | 0.341435 | 1 | 0.004472 | 0.004472 | 6.993958 | 0.010638 | Comamonas_day3 |
| I:P:C | 0.056775 | 0.046169 | 1.229724 | 0.224035 | 1 | 0.000896 | 0.000896 | 1.401764 | 0.24152 | Comamonas_day3 |
| I:P:A | 0.064206 | 0.047309 | 1.357161 | 0.180274 | 1 | 0.003262 | 0.003262 | 5.101258 | 0.027895 | Comamonas_day3 |
| I:C:A | 0.079412 | 0.048423 | 1.639966 | 0.10672 | 1 | 0.002153 | 0.002153 | 3.367361 | 0.07191 | Comamonas_day3 |
| P:C:A | 0.019028 | 0.047309 | 0.402206 | 0.689091 | 1 | 0.00137 | 0.00137 | 2.142584 | 0.148953 | Comamonas_day3 |
| I:P:E | -0.03737 | 0.050576 | -0.73881 | 0.463166 | 1 | 0.002353 | 0.002353 | 3.679592 | 0.060279 | Comamonas_day3 |
| I:C:E | 0.075164 | 0.047309 | 1.58877 | 0.117845 | 1 | 0.000569 | 0.000569 | 0.889196 | 0.349819 | Comamonas_day3 |
| P:C:E | 0.013183 | 0.051619 | 0.255393 | 0.799372 | 1 | 0.001026 | 0.001026 | 1.603982 | 0.210677 | Comamonas_day3 |
| I:A:E | 0.028097 | 0.048423 | 0.58025 | 0.564116 | 1 | 1.15E-05 | 1.15E-05 | 0.018027 | 0.893685 | Comamonas_day3 |
| P:A:E | -0.0561 | 0.051619 | -1.08679 | 0.281871 | 1 | 0.000493 | 0.000493 | 0.771344 | 0.383622 | Comamonas_day3 |
| C:A:E | 0.016121 | 0.050576 | 0.318754 | 0.751121 | 1 | 9.37E-05 | 9.37E-05 | 0.146473 | 0.703404 | Comamonas_day3 |
| I:P:C:A | 0.000148 | 0.06364 | 0.002332 | 0.998148 | 1 | 0.000243 | 0.000243 | 0.380685 | 0.539783 | Comamonas_day3 |
| I:P:C:E | -0.00973 | 0.066104 | -0.14726 | 0.883469 | 1 | 0.000976 | 0.000976 | 1.526947 | 0.221823 | Comamonas_day3 |
| I:P:A:E | 0.032286 | 0.066906 | 0.482566 | 0.631319 | 1 | 4.69E-05 | 4.69E-05 | 0.073273 | 0.787642 | Comamonas_day3 |
| I:C:A:E | -0.04466 | 0.066104 | -0.6756 | 0.502126 | 1 | 0.002298 | 0.002298 | 3.593933 | 0.06325 | Comamonas_day3 |
| P:C:A:E | 0.075936 | 0.06848 | 1.108874 | 0.27231 | 1 | 0.000347 | 0.000347 | 0.542535 | 0.464514 | Comamonas_day3 |
| I:P:C:A:E | -0.0751 | 0.090591 | -0.82901 | 0.41068 | 1 | 0.000439 | 0.000439 | 0.687264 | 0.41068 | Comamonas_day3 |
| I | 0.016893 | 0.022414 | 0.753652 | 0.453866 | 1 | 0.18672 | 0.18672 | 247.7677 | 1.64E-23 | Comamonas_day11 |
| P | 0.004612 | 0.022414 | 0.20576 | 0.837642 | 1 | 0.439429 | 0.439429 | 583.0991 | 1.5E-33 | Comamonas_day11 |
| C | 0.213397 | 0.02506 | 8.515412 | 4.49E-12 | 1 | 2.63E-05 | 2.63E-05 | 0.03491 | 0.852385 | Comamonas_day11 |
| A | 0.359452 | 0.022414 | 16.03665 | 6.4E-24 | 1 | 0.073469 | 0.073469 | 97.4897 | 2.05E-14 | Comamonas_day11 |
| E | 0.063809 | 0.022414 | 2.84678 | 0.005954 | 1 | 3.66E-05 | 3.66E-05 | 0.048573 | 0.826277 | Comamonas_day11 |
| I:P | -0.03231 | 0.031699 | -1.0192 | 0.312008 | 1 | 0.158625 | 0.158625 | 210.4872 | 9.36E-22 | Comamonas_day11 |
| I:C | -0.19826 | 0.033622 | -5.89694 | 1.59E-07 | 1 | 8.21E-05 | 8.21E-05 | 0.108913 | 0.74248 | Comamonas_day11 |
| P:C | -0.22961 | 0.033622 | -6.82916 | 3.96E-09 | 1 | 0.000408 | 0.000408 | 0.541872 | 0.46439 | Comamonas_day11 |
| I:A | -0.27124 | 0.031699 | -8.55672 | 3.81E-12 | 1 | 0.034609 | 0.034609 | 45.92374 | 4.89E-09 | Comamonas_day11 |
| P:A | -0.37205 | 0.031699 | -11.737 | 1.68E-17 | 1 | 0.096147 | 0.096147 | 127.5813 | 8.73E-17 | Comamonas_day11 |
| C:A | -0.33551 | 0.033622 | -9.97899 | 1.36E-14 | 1 | 0.040735 | 0.040735 | 54.05267 | 4.85E-10 | Comamonas_day11 |
| I:E | -0.07342 | 0.031699 | -2.31632 | 0.023807 | 1 | 0.005326 | 0.005326 | 7.066753 | 0.009942 | Comamonas_day11 |
| P:E | -0.06824 | 0.031699 | -2.15262 | 0.035186 | 1 | 0.00017 | 0.00017 | 0.225734 | 0.636348 | Comamonas_day11 |
| C:E | -0.08561 | 0.033622 | -2.54617 | 0.013349 | 1 | 0.000503 | 0.000503 | 0.667726 | 0.416925 | Comamonas_day11 |
| A:E | -0.01183 | 0.031699 | -0.37327 | 0.710202 | 1 | 0.002069 | 0.002069 | 2.745127 | 0.102524 | Comamonas_day11 |
| I:P:C | 0.20761 | 0.046209 | 4.492897 | 3.07E-05 | 1 | 9.15E-06 | 9.15E-06 | 0.012146 | 0.912594 | Comamonas_day11 |
| I:P:A | 0.279414 | 0.044829 | 6.232912 | 4.25E-08 | 1 | 0.038854 | 0.038854 | 51.55752 | 9.68E-10 | Comamonas_day11 |
| I:C:A | 0.289976 | 0.046209 | 6.27538 | 3.59E-08 | 1 | 0.028995 | 0.028995 | 38.47479 | 4.78E-08 | Comamonas_day11 |
| P:C:A | 0.349205 | 0.046209 | 7.557146 | 2.12E-10 | 1 | 0.047598 | 0.047598 | 63.15989 | 4.41E-11 | Comamonas_day11 |
| I:P:E | 0.084312 | 0.044829 | 1.88076 | 0.064628 | 1 | 0.005672 | 0.005672 | 7.526915 | 0.007907 | Comamonas_day11 |
| I:C:E | 0.085668 | 0.046209 | 1.853952 | 0.068429 | 1 | 0.001914 | 0.001914 | 2.539442 | 0.116039 | Comamonas_day11 |
| P:C:E | 0.102346 | 0.046209 | 2.214865 | 0.030394 | 1 | 0.00141 | 0.00141 | 1.870794 | 0.176244 | Comamonas_day11 |
| I:A:E | -0.05384 | 0.044829 | -1.20093 | 0.234275 | 1 | 0.000723 | 0.000723 | 0.959532 | 0.331051 | Comamonas_day11 |
| P:A:E | 0.011823 | 0.044829 | 0.263736 | 0.792844 | 1 | 0.000694 | 0.000694 | 0.921301 | 0.340803 | Comamonas_day11 |
| C:A:E | 0.024505 | 0.046209 | 0.53031 | 0.597761 | 1 | 0.000106 | 0.000106 | 0.141264 | 0.70829 | Comamonas_day11 |
| I:P:C:A | -0.30119 | 0.064381 | -4.67829 | 1.58E-05 | 1 | 0.031511 | 0.031511 | 41.81333 | 1.69E-08 | Comamonas_day11 |
| I:P:C:E | -0.10451 | 0.064381 | -1.62329 | 0.109522 | 1 | 0.003455 | 0.003455 | 4.584639 | 0.036135 | Comamonas_day11 |
| I:P:A:E | 0.044263 | 0.063398 | 0.69818 | 0.487634 | 1 | 0.000945 | 0.000945 | 1.254367 | 0.266972 | Comamonas_day11 |
| I:C:A:E | 0.006165 | 0.064381 | 0.095763 | 0.924013 | 1 | 6.67E-05 | 6.67E-05 | 0.08855 | 0.767008 | Comamonas_day11 |
| P:C:A:E | -0.03821 | 0.064381 | -0.59347 | 0.554991 | 1 | 0.000364 | 0.000364 | 0.48345 | 0.48942 | Comamonas_day11 |
| I:P:C:A:E | 0.013393 | 0.090355 | 0.148222 | 0.882641 | 1 | 1.66E-05 | 1.66E-05 | 0.02197 | 0.882641 | Comamonas_day11 |
